# Automatic Individual Cortical Parcellation for the Human Connectome Project

**DOI:** 10.1101/2025.04.29.651219

**Authors:** Chunhui Yang, Timothy S. Coalson, Seyedeh-Rezvan Farahibozorg, Janine D. Bijsterbosch, Stephen M. Smith, David C. Van Essen, Matthew F. Glasser

## Abstract

The original Human Connectome Project multimodal cortical parcellation (HCP_MMP1.0) used MRI-derived local features and long-distance functional connectivity measures to define a multimodal parcellation at the group level, accompanied by an automated areal classifier, to create subject-specific mappings of the human cerebral cortex. These mappings, referred to as individual (cortex) parcellations, aim to capture individual variability in areal organization by learning from both structural and functional data. However, a strict supervised learning approach using the group parcellation as labels would have no incentive to learn individual differences that registration is unable to reconcile (e.g., atypical 55b topologies). Furthermore, there are many types of resting state network (RSN) feature maps, and it is unclear which type would most accurately or effectively classify areas, or even what should be the primary criteria for evaluating classification performance. Here, we introduce an <u>A</u>real <u>R</u>ecognition <u>E</u>nsemble with <u>N</u>ested <u>A</u>pproach (ARENA) classifier that learns from uncertain labels, using a novel application of weakly supervised learning to this type of problem. Additionally, in comparing multiple candidate RSN decompositions, temporal ICA and PROFUMO maps outperformed the original spatial ICA-based approach based on objective criteria. With these refinements, the ensemble classifier achieved a reliable individual variability score of 8380, an average areal detection rate of 97.8%, and test-retest reproducibility of 73.3%, outperforming a retrained version of the original Multi-layer Perceptron (MLP) model (whose reliable individual variability score was 3128, average areal detection rate was 97.2%, and test-retest reproducibility was 71.6% on the same dataset). Furthermore, the ARENA classifier demonstrated stronger generalization for all three measures when applied to task fMRI data that were not part of the training dataset. Using the refined classifier and leveraging all 1071 HCP-Young Adult subjects, we identified new types of atypical organization of language-related area 55b. Here we provide the fully data-driven HCP_MMP1.0_1071_MPM (Maximum Probability Map) group parcellation and a summary of area 55b organization in both hemispheres. Our automated individual parcellation pipeline powered by the novel ARENA classifier is now integrated into the HCP pipelines, offering a user-friendly tool for the neuroimaging community.

## 1. Introduction

Cerebral cortex is a thin sheet of tissue that can be subdivided into a mosaic of cortical areas - regions that differ reliably from neighboring areas in one or more of four basic categories: function, architecture, connectivity, and/or topographic organization (Van Essen & Glasser, 2018). Classical anatomists debated whether human cortex contains only ∼50 areas as suggested by cytoarchitecture (Brodmann, 1909) or perhaps as many as ∼200 areas as suggested by myeloarchitecture (Nieuwenhuys et al., 2015; Vogt & Vogt, 1919). The advent of noninvasive MRI late in the 20^th^ century provided many additional modalities for identifying cortical areas in individual and group averaged data. These include identification of areas based on the T1w/T2w ratio, a non-invasive measure of greymatter myelin content, with areal boundaries identified by spatial gradients (Glasser & Van Essen, 2011) that agree with quantitative postmortem cytoarchitecture (Fischl et al., 2008). Many investigators have reported surface-based parcellations using only resting-state fMRI correlations, or ‘functional connectivity’ (Gordon et al., 2016; Power et al., 2011; Schaefer et al., 2018; Yeo et al., 2011). However, many constituent parcels do not correspond to well-defined topographically organized early somatosensory, motor, and visual areas (Van Essen & Glasser, 2018).

Major progress in comprehensively mapping human cortical areas has come from a multimodal, semi-automated, supervised, gradient-based parcellation that identified 180 areas in each hemisphere using hundreds of subjects from the young adult HCP (Glasser et al., 2016) and using modalities related to function (task-fMRI activation patterns), architecture (cortical thickness and T1w/T2w-based myelin), connectivity (from resting-state fMRI), and topography (also derived from resting-state fMRI). Accurate parcellation was highly dependent on precise surface-based alignment of individuals to a group average using areal-feature-based alignment (Robinson et al., 2018; Robinson et al., 2014). An areal classifier based on a multi-layer perceptron (MLP) successfully identified nearly all areas in nearly all 210 subjects used to generate the original group average and train the classifier (the ‘210P’ group) and in another group of 210 subjects used for ‘validation’/evaluation (the ‘210V’ group). Detailed studies of one area, language-related area 55b, revealed that it had a distinctive topology in about 10% of individuals, being either ‘split’ into two parts by an intervening area or ‘switched’ (previously called ‘shifted’) dorsally, switching places with neighboring area FEF.

Classical anatomical studies have shown that well-defined areas such as V1 differ in size (surface area) by two-fold or more across individuals (Andrews et al., 1997; Stensaas et al., 1974). Areal boundaries also vary across individuals in relation to cortical folds in humans, but much less so in macaques (Hilgetag & Barbas, 2006; Van Essen, 1997; Van Essen et al., 2019). Furthermore, inter-individual variability in brain anatomy and parcellation can predict aspects of individual behavior, such as cognitive performance and personality traits (Bijsterbosch et al., 2018; Cachia et al., 2018; Kong et al., 2019; Kong et al., 2021). Achieving robust, reproducible and reliable individual parcellations of human cerebral cortex is vital for attaining a deeper understanding of the nature of individual variability. Such an endeavor serves as the foundational goal behind the development of the areal classifier. (Otherwise, simply imposing a group parcellation atlas on each individual would be sufficient.) The areal classifier is specifically designed to accurately represent individual variability that is not already captured by areal-feature-based registration, thereby enabling a deeper understanding of its functional significance.

The MLP areal classifier used by Glasser et al. (2016) was initially trained using all datasets to select a subset of subjects having an area well-matched to the group average for subsequent training of that area’s areal classifier. Important features distinguishing each area were identified during this first stage of training through derivatives obtained by calculating the gradients of the MLP’s output with respect to the input nodes. This subject filtering stage facilitated the reasonable utilization of the group parcellation as training labels for the second stage. This method treated HCP_MMP1.0 as ground truth labels during training, partially neglecting the uncertainty inherent in group parcellation when applied to individual subjects. (This “partial neglect” was acknowledged, as this method incorporated a preprocessing stage to select only subjects with typical area topology and good contrast for final classifier training.) An attractive alternative approach is to use weakly supervised learning (Zhou, 2017), which explicitly considers label uncertainty during training and does not exclude valuable data from poorly matching subjects.

The previous study (Glasser et al., 2016) reported measures such as averaged areal detection rate and test-retest reproducibility to describe the overall performance of the areal classifier. However, these measures would be artificially high if areal classifiers learned to perfectly predict the group training labels. Thus, we must consider carefully the purpose of the areal classifier—to identify reproducible individual variability in human cortical areas and develop measures that directly assess performance on this task. This motivates an evaluation framework that focuses on the following two aspects: 1) valid reproducibility (i.e., the ability to capture reproducible individual variability differing from the group average) and 2) generalization ability (i.e., the ability to make reasonable predictions on unseen types of fMRI data, e.g., task fMRI). This framework directly addresses the purpose of the areal classifier and makes it harder for aberrant classifiers to “game the system” (e.g., classifiers that memorize the locations of group training labels instead of learning multi-modal areal fingerprints as intended).

In this paper, we introduce an Areal Recognition Ensemble with Nested Approach (ARENA) classifier. In the first stage, a XGBoost ensemble model combines predictions from multiple base learners to generate individual areal probabilities. Subsequently, a regression model is employed to learn from those predicted probabilities, explicitly taking into account label uncertainty. ARENA constitutes a weakly supervised learning model designed to address label uncertainty stemming from using the group parcellation HCP_MMP1.0 at the individual level. In the initial stage, each base learner learns from partial in- and outside-areas within the search region, collectively contributing to the initial prediction. Then the regression model uses these initial predictions to directly learn the uncertainty. Additional methodological explorations (see methods section Resting State Network (RSN) features for a detailed discussion) in this manuscript consider two alternatives to spatial ICA maps as topographical features, namely temporal ICA maps (Glasser et al., 2018, 2019) and Probabilistic Functional Mode (PFM) maps (Farahibozorg et al., 2021; Harrison et al., 2020; Harrison et al., 2015). We compare the performance of all feature sets and use the refined areal classifier to determine the optimal RSN decomposition types.

We also develop an evaluation framework using previously developed and newly introduced metrics that consider 1) reliability, the reproducibility of individual variability in test-retest subjects, 2) robustness, the similarity between predicted group-level parcellation and the HCP_MMP1.0, and 3) generalizability, the method’s ability to generalize across unseen subjects and fMRI paradigms. Given these criteria, we demonstrate that the refined areal classifier substantially outperforms the existing MLP.

Using this refined areal classifier, we analyze area 55b of 1071 3T subjects from the HCP-Young Adult cohort. We confirm the presence of two previously reported topological patterns (‘split’ and ‘switched,’ previously called ‘shifted’) and demonstrate two new types of atypical topology. A fully automated, data-driven refined group parcellation HCP_MMP1.0_1071_MPM (Max Probability Map) is produced as a refinement to the original HCP_MMP1.0, yielding a parcellation that we consider closer to ground truth.

## 2. Methods

### 2.1 Dataset information

#### Participants

We used multimodal data from the HCP Young Adult (HCP-YA) project, acquired in compliance with the Washington University Institutional Review Board (Van Essen et al., 2013). These data include 1071 HCP-YA participants (ages 22 – 35) scanned at 3T (45 test-retest subjects were scanned twice) who had successfully completed 3T preprocessing by the HCP Pipelines (Glasser et al., 2013) refined as described below. The resting-state fMRI and task-fMRI data included spatial denoising using sICA+FIX (Salimi-Khorshidi et al., 2014), spatial ICA recleaning (Yang et al., 2024), areal-feature-based surface registration using MSMAll, and temporal ICA denoising (Glasser et al., 2018, 2019). The MSMAll algorithm was based on the original angular deviation model with higher order regularization (Glasser et al., 2016; Robinson et al., 2014), not the strain-based MSM regularization model of Robinson et al. (2018).

#### HCP-YA images

T1w and T2w HCP-YA structural scans were described previously (Glasser et al., 2022; Glasser et al., 2013; Ugurbil et al., 2013). The 3T resting state fMRI data were acquired with 2.0 mm isotropic resolution, TR=720 ms and 1200 frames (14.4 min) for each of 4 runs. Two runs each were acquired with reversed phase encoding directions (RL or LR) with the order counterbalanced across each of two scan sessions (Smith et al., 2013) for a total of 4800 frames of resting state per subject along with matching phase reversed spin echo fMRI scans for distortion correction. The 3T task fMRI data (Barch et al., 2013) were acquired using identical pulse sequence settings while participants performed 7 tasks (working memory, gambling, motor, language, social cognition, relational processing and emotion processing) with runs lasting between 2 and 5 min (176-405 frames) and totaling 22.6 min (1884 frames) in session 1 and 24 min (1996 frames) in session 2 for a total of 3880 frames of task fMRI per subject. Each task scan comprised a pair of runs with reversed phase encoding directions, first RL then LR. Runs were halted if subjects stopped performing any task.

#### Image preprocessing

All datasets were preprocessed using the HCP Pipelines (Glasser et al., 2013), including three structural spatial preprocessing pipelines, two functional preprocessing pipelines, a multi-run spatial ICA+FIX fMRI artifact removal pipeline (Salimi-Khorshidi et al., 2014; Glasser et al., 2018, 2019) plus a ‘recleaning’ process to further improve classification of sICA components (Yang et al., 2024), the multi-modal ‘MSMAll’ areal-feature-based surface registration (Glasser et al., 2016; Robinson et al., 2014), and the temporal ICA fMRI artifact and nuisance removal was applied to the full group (Glasser et al., 2018, 2019). Cortical surface data were represented on a 32k standard ‘grayordinates’ mesh for each hemisphere’s cortical surface (Glasser et al., 2013). The average vertex spacing on individual midthickness surfaces is ∼2 mm, but the spacing varies across regions.

### 2.2 Individual areal classifier

#### Classification problem

The semi-automated gradient-based multimodal parcellation method introduced by Glasser et al. (2016) identified 180 cortical areas in each hemisphere based on the HCP 210P dataset, resulting in the HCP_MMP1.0 parcellation. However, an analogous gradient-based approach would be inordinately time-consuming to perform on individual subjects and might perform poorly owing to low signal-to-noise within individuals. To address this challenge, our previous study used a multi-layer perceptron designed to learn the unique multimodal characteristics (areal fingerprint) of each cortical area, distinguishing it from the surrounding cortex. This study follows the same binary classification approach for each area, which can be summarized as follows:

For each parcel labeled according to the HCP_MMP1.0 parcellation, a searchlight with a radius of 30 mm (geodesic distance – on the individual’s midthickness surface) was established to set a reasonable spatial limit on candidate vertices to be classified. As described in Glasser et al., 2016, 30 mm represents a tradeoff between missing individual variability and avoiding geographically remote areas that may have a similar areal fingerprint (e.g., because they are within the same functional network). Subsequently, a binary classifier was trained using feature vectors from all vertices within the searchlight region. The classifier was tasked with distinguishing between ‘inside-area’ positive labels (corresponding to vertices inside the labeled area from HCP_MMP1.0 parcellation) and ‘outside-area’ negative labels (vertices outside the labeled area from the HCP_MMP1.0 parcellation). A total of 360 classifiers were trained to predict whether a given vertex was inside an area or not across both hemispheres of the whole brain.

To illustrate this process, Fig 2 shows group-average areas 55b and RSC (colored fuchsia) from the training dataset along with the associated searchlight regions (yellow). Collectively, these regions form the overall search space.

#### Individual parcellation pipeline

The pipeline aims to produce individual parcellations with pretrained model weights, as illustrated in Fig 1B. It begins by creating the required multimodal feature maps (discussed in detail below) for a specific subject and then uses the 360 trained areal classifiers for inference. The resulting probabilistic maps of each area are merged by selecting the top probability for each vertex across maps. The resulting parcellation undergoes regularization through a series of post-processing steps (Glasser et al., 2016): 1) removing patches smaller than 25 mm² on the individual’s midthickness surface, 2) connecting disjoint areas within a few vertices of another area, and 3) ensuring that the remaining pieces were at least 0.33 times the size of the largest piece and within 30 mm of its nearest edge.

**Figure 1.**
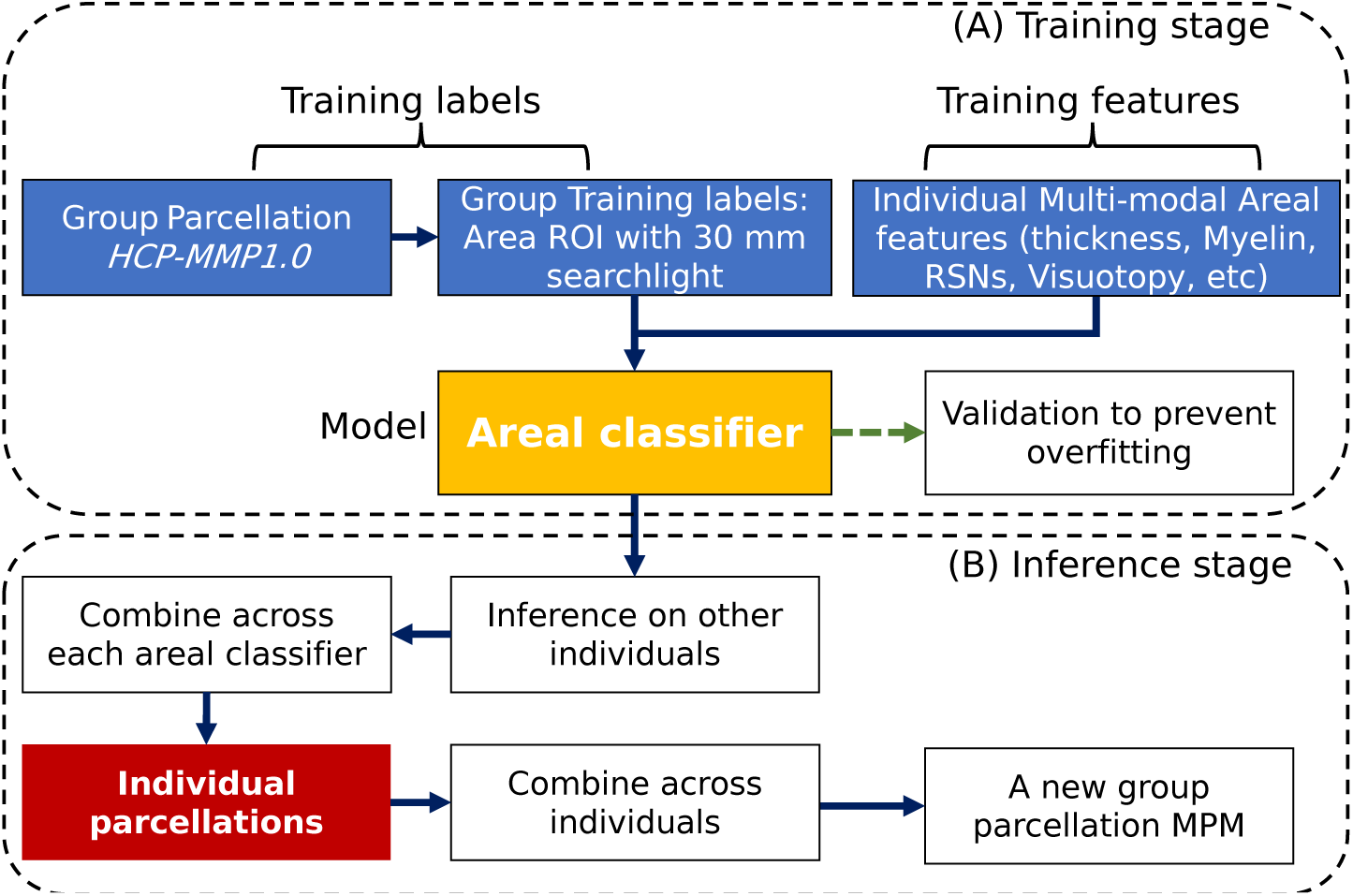
The individual parcellation framework. (A) The training stage including training labels (HCP_MMP1.0_210P_Orig in this study) from the original version 1 of the HCP’s multi-modal parcellation, multimodal training features and an areal classifier model that can be replaced by advanced machine learning algorithms. (B) The inference stage to derive individual parcellations and a new group parcellation on a group of subjects. This stage can be regarded as the individual parcellation pipeline with trained model weights.

**Figure 2.**
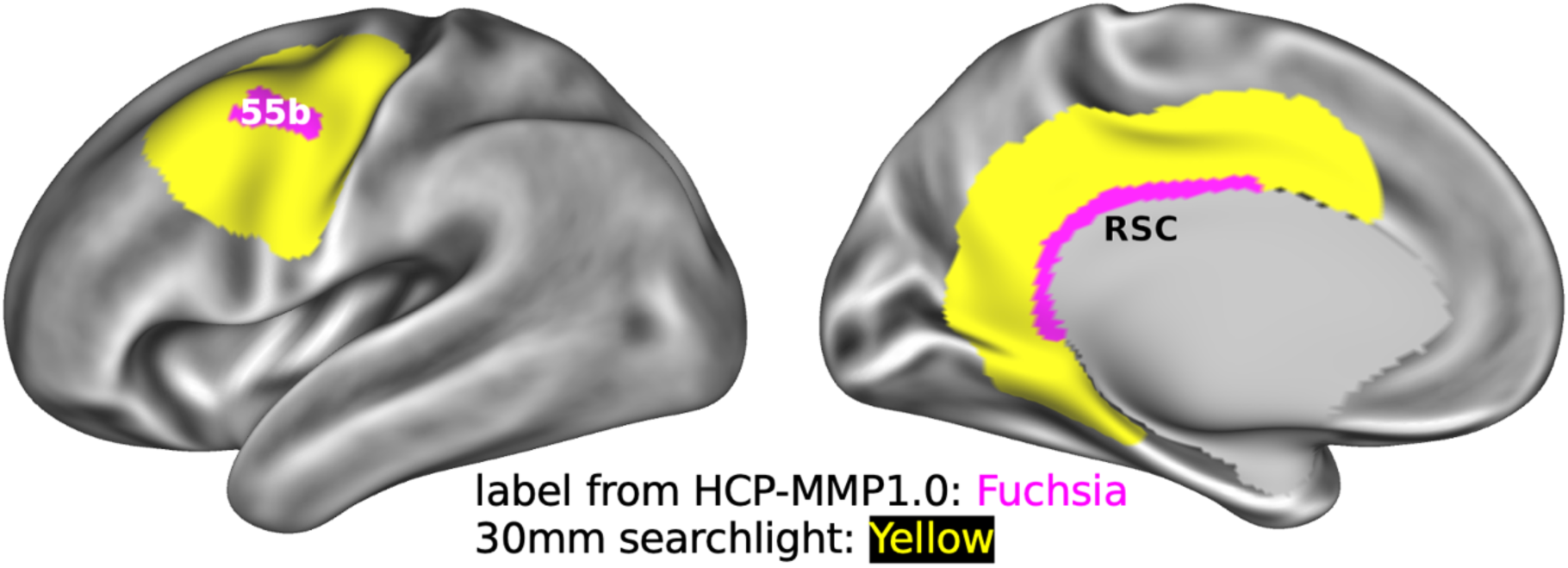
Overview of the classification problem. Classification problem for 55b and RSC (an example to show that the medial wall is excluded automatically and is never part of the searchlight) in the left hemisphere with a 30mm searchlight indicated by yellow regions around the HCP_MMP1.0 labels (fuchsia; on top).

#### Multimodal features

We adopted a set of individual-level feature maps (Glasser et al., 2016) of cortical thicknesses (with curvatures regressed out), surface curvatures, T1w/T2w ratios (myelin maps), vein effects, fMRI dropouts, insular surface artifacts, resting-state timeseries standard deviations, regression coefficients of mean gray time courses (beta values), and topographic regression dot products that aid in identifying visuotopically organized areas. Additionally, we introduced multiple types of carefully designed resting state network (RSN) maps (different from the previous approach), as described in the next section.

#### Resting State Network (RSN) features

RSN feature maps are the most informative single modality for areal classification (Glasser et al., 2016). We explored five distinct sets of RSN feature maps, all based on tICA cleaned HCP datasets, that differ in how they were generated and what they represent:

1. Individualized Spatial ICA Maps (sICA): The individual spatial ICA maps were obtained following the first three steps from the tICA pipeline (Yang et al., 2024): MIGP (Melodic’s Incremental Group PCA), group spatial ICA, and weighted multiple regression (Glasser et al., 2016) with temporal ICA cleaned HCP 3T 1071 resting state fMRI dense timeseries as the input. Two group spatial ICA decompositions were derived after fitting five (resulting dimensionality: d=92) and six (d=76) Wishart distributions; each map was manually inspected by MFG and either included robust cortical features (n=55 for d=92; n=48 for d=76) or were dominated by non-cortical features (n=37 for d=92; n=28 for d=76). These individual spatial ICA maps were generated from the 1071 dataset and have multiple bugfixes in the relevant pipeline relative to the original d=137 from HCP 210P (Glasser et al., 2016). Importantly, spatial ICA maps are constrained to be spatially uncorrelated with each other.
2. Individualized Temporal ICA Spatial Maps (tICA): These individual temporal ICA maps were derived after obtaining the above individual sICA maps following the subsequent steps from the tICA pipeline: concatenation of individual sICA timeseries, group temporal ICA and single temporal regression of the unconcatenated individual tICA time courses into each individual’s fMRI data. These recently introduced maps (Glasser et al., 2018, 2019; Yang et al., 2024) enhance the feature space by providing higher Signal-to-Noise Ratio (SNR) features (extra signal is embedded in the tICA spatial maps by including the connectivity between functional networks while noise levels stay the same). A major difference between individual tICA maps and individual sICA maps is that, for temporal ICA, the connectivity information resides in the spatial maps (because the temporal ICA timeseries components are independent), whereas for spatial ICA, the connectivity information resides in both the sICA maps and the temporal correlations between component timeseries (i.e., within the network matrices). No manual selection was conducted, and all components were used as features (the same d=92 and d=76 as above). Temporal ICA decompositions can be extended to new individuals and datasets by first performing weighted multiple regression using the same starting sICA decomposition and then remixing the components according to the temporal ICA mixing matrix.
3. Individualized PROFUMO original “noise-free spatial maps” (PFM original maps; origmaps): The PROFUMO method simultaneously estimates subject and group probabilistic functional mode (PFM) maps and network matrices, instead of separate parcellation and mapping steps (Farahibozorg et al., 2021; Harrison et al., 2020; Harrison et al., 2015). The signal-only PFM original spatial maps (i.e., with noise removed) were used as PFM-origmaps for subsequent analysis. The sessions from the 1071 subjects and 45 retest subjects were analyzed via the PROFUMO algorithm using the dimensionalities of d=92 and d=76 PFMs to match the dimensionalities above from group spatial and temporal ICA. PFM timeseries were constrained by the Hemodynamic Response Function (HRF) and, for each subject, a single temporal network matrix, and a single set of spatial maps were estimated across all runs, both informed by group-level priors. The default correction for the spatial smoothness of the data was applied with a value of 0.5. To reduce computational demands, the ranks 5*92=460 (d=92) and 5*76=380 (d=76) of the SVD-based approximation were used. Importantly, PFMs are not constrained to be either spatially or temporally independent of each other, but rather must all have the properties of BOLD functional networks.
4. Individualized PROFUMO spatial maps after dual regression (PFM-DR): A variant of PROFUMO spatial maps was designed to revert the spatial denoising performed by PROFUMO and be able to apply the PFM decompositions to new spaces such as volume space. Once the original PROFUMO spatial maps for each individual were obtained, they were regressed into each individual’s dense timeseries to obtain the corresponding timeseries (stage 1). These were subsequently regressed back into the same dense timeseries, yielding a set of PROFUMO spatial maps (stage 2). This approach maintains the decomposition of PROFUMO, but removes the spatial map denoising (reintroduces the noise and reduces the contrast in vs outside the functional network). In practice, we expect this approach to be similar to using the “noisy PFM spatial maps” that are produced in the PROFUMO framework, except that the dual regression approach makes the mathematics of “noise/residual” estimation more similar to individualized sICA/tICA, thus providing a fairer comparison of “signal” estimation across methods. Importantly, PFM-DR requires the individual subject PFM origmaps to exist as the starting input, so this method cannot currently be easily generalized to subjects that were not part of the original PFM decomposition.
5. Individualized PROFUMO Spatial Maps obtained by Weighted Regression (PFM-WR): Unlike PFM original maps (PFM-origmaps, #3) and PFM dual regression maps (PFM-DR, #4), which cannot currently be easily extended to unseen subjects or fMRI runs, this approach serves as a workaround solution. In this method, the group decomposition from PROFUMO is regressed into the individual fMRI matrix using the same weighted multiple regression process as in sICA. This approach enables the generation of individual PFM weighted resting-state network (RSN) spatial maps and their corresponding timeseries, although it loses the benefits of the concurrent group and individual estimation in PROFUMO.

The Results section extensively compares the performance of each of these RSN map representations, including their ability to capture reproducible individual variability differing from the group parcellation and the generalizability of each approach.

#### Dataset groups

The primary data groups involve splitting the HCP 3T 1071 group with resting state fMRI, with the following allocations (also see Table 1): 1) training dataset with 475 subjects, called the ‘475P’ group by analogy to the 210P (‘parcellation’ training dataset) used by Glasser et al. (2016); 2) validation dataset with 50 subjects, called the ‘50T’ group by analogy to the 29T (test dataset, used to prevent overfitting) used by Glasser et al. (2016); and 3) main evaluation datasets (Eval #1-R) with 475 subjects, called ‘475V’ by analogy to the 210V (final validation dataset) used by Glasser et al. (2016). In addition to these main data groups, we set aside the 45 test-retest subjects to create two pairs of independent evaluation datasets (Eval test-R and Eval retest-R) that were crucial for assessing the reproducibility and reliability of each model. To gauge the model’s ability to generalize to fMRI data beyond resting-state scans, we also incorporated task-based fMRI from the 475V group and the test-retest group as additional evaluation datasets (Eval #1-T, Eval test-T and Eval retest-T). HCP-YA families were always grouped together to avoid leakage of genetically-based information across groups.

**Table 1.** Overview of the dataset groups.

| Dataset name | HCP dataset | # session | fMRI data |
| --- | --- | --- | --- |
| Train (475P) | Young Adult | 475 | Resting state |
| Validation (50T) | Young Adult | 50 | Resting state |
| Evaluation (475V; Eval #1) | Young Adult | 475 | Resting state (R), Task fMRI (T) |
| Evaluation (45V; Eval test-T) | Young Adult | 45 | Resting state, Task fMRI |
| Evaluation (45V; Eval retest-T) | Young Adult | 45 | Resting state, Task fMRI |

#### Baseline models

The baseline model adheres to the same classification framework outlined in the *Classification problem* subsection. The model is the original Multi-Layer Perceptron (MLP) implemented in MATLAB (Glasser et al., 2016), which was initially designed for classifying resting-state networks (Hacker et al., 2013). Its main architecture includes an MLP with one hidden layer containing 9 nodes. The training process involved an initial stage aimed at identifying important features and selecting a subset (approximately 75%) of well-aligned subjects for each area. Subsequently, a final training stage was conducted on the well-aligned subjects with all features for each area.

#### Data normalization

Data normalization is crucial in machine learning to ensure that features have consistent scales, enabling algorithms to converge faster, perform better, and be more robust to outliers, ultimately improving model accuracy and interpretability. We re-used the data normalization strategy introduced in the MLP (Glasser et al., 2016): Feature-wise and Type-wise Normalization. Within each area’s search space, features were normalized by subtracting the mean of each *feature* per individual within the search space and dividing by the median standard deviation for each feature type per individual within the search space.

#### Label uncertainty

Label uncertainty arises from two main factors. Firstly, it stems from the use of a particular group parcellation, such as HCP_MMP1.0, which represents the best available estimate for defining multi-modal cortical areas but may be improved in the future. Secondly, label uncertainty is exacerbated by inter-subject variability that cannot be represented as a smooth registration deformation (topological variants). In the classification problem, the group parcellation serves as the label for every subject in the training group, ignoring inherent inter-subject variability. The MLP model excluded ∼25% of subjects having poor feature map alignment during the training phase, as it presumes that models can learn correct areal features from subjects having typical areal organization. Because the MLP model does not effectively consider label uncertainty during training, it may show suboptimal generalization performance. To address this challenge, we reframed the classification problem as a form of weakly supervised learning, specifically, inaccurate supervision (Zhou, 2017). Our approach assumes the presence of noisy labels that can provide supervision signals within a supervised learning framework. It leverages a combination of ensemble learning, a straightforward yet highly effective and widely used method (Mordelet & Vert, 2014; Sheng et al., 2008; Snow et al., 2008; Zhou, 2012), and a dual-stage training schema that enables direct learning on the probabilities generated by the initially trained model, akin to the commonly used strategy of Pseudo-labeling in the context of semi-supervised learning (Cascante-Bonilla et al., 2021; Rizve et al., 2021).

Specifically, we employed ensemble learning via a bagging (bootstrap aggregating) strategy: we trained 100 XGBoost classifiers in the first stage and another 100 XGBoost regressors in the second stage, each on a randomly selected subset of the training samples (70% of the training samples without replacement). This approach introduces diversity across base learners and reduces variance, improving the model’s robustness to label noise and subject-level variability. The final prediction of stage #1 or stage #2 is obtained by averaging the class probabilities across all base classifiers. Hyperparameters for the XGBoost learners (e.g., maximum tree depth, learning rate, and number of boosting rounds) were tuned using five-fold cross-validation on the training data, and early stopping was applied based on validation performance.

#### Model architecture

We chose an ensemble of XGBoost models in our architecture for several reasons. XGBoost (Fig 3E), a gradient boosting algorithm that sequentially constructs decision trees where each tree is trained to correct the previous one’s errors, and individual tree predictions are combined by summing them to produce the final probability, has exhibited robustness to outliers and noise, reducing the need for fine-tuning of hyperparameters (Chen & Guestrin, 2016). Furthermore, these models permit combining multiple ‘weak learners’ within our ensemble for weakly supervised learning (Claesen et al., 2015; Høie et al., 2023; Zhao et al., 2021) ultimately yielding a robust final prediction.

**Figure. 3.**
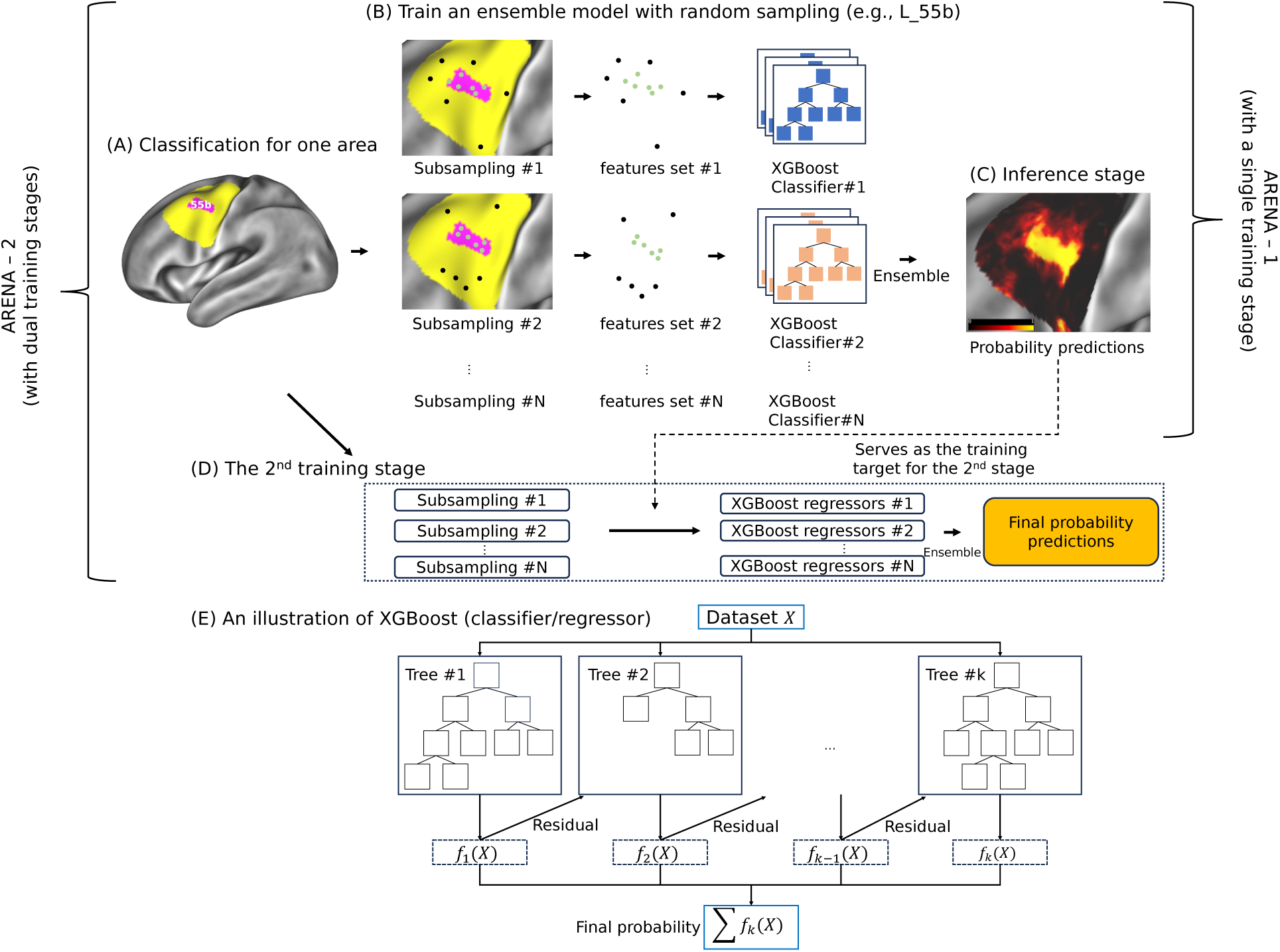
Overview of the individual parcellation method under a weakly supervised setting. (A) Classification problem for 55b in the left hemisphere with a 30 mm searchlight indicated by yellow regions around the HCP_MMP1.0 labels (fuchsia; on top). (B) ARENA-1 model for weakly supervised learning (N stands for the number of XGBoost learners). (C) Inference stage with final predictions obtained by averaging base learner predictions. (D) The 2^nd^ training stage then considers the label uncertainty from the 1^st^ stage using the predicted probabilities as training targets. Combining ARENA-1 and the 2^nd^ stage results in the ARENA-2 model. (E) A high-level illustration of the XGBoost model. XGBoost is a sequential ensemble learning algorithm that builds an ensemble of decision trees to improve predictions incrementally. It combines individual tree predictions by summing them to produce the final probability or prediction.

Beyond incorporating a model that addresses label uncertainty, we designed a dual-stage training process for the areal classifier to further mitigate the impact of label uncertainty. While the ensemble of XGBoost classifiers in the first round helps reduce the influence of noisy labels by averaging predictions across diverse subsets of training data, it does not explicitly model or correct underlying noise in the labels themselves. Therefore, we introduced a second stage that treats the ensemble’s averaged soft predictions as a form of probabilistic supervision, enabling models to learn from the aggregated uncertainty rather than relying solely on the original (potentially inaccurate) hard labels. The first training stage (ARENA-1; Fig 3B & C) is responsible for generating coarse areal probabilities, while the second stage employs a regression model targeting these probabilities (Fig 3D). ARENA-2 is defined as an integrated model with ARENA-1 plus the 2^nd^ training stage. In the first stage, the classifier comprises 100 XGBoost base learners (classifiers) for each area. The probability predictions from each base learner are averaged to yield the ultimate prediction. The hyperparameters for these XGBoost base learners were optimized manually using the 50T validation dataset. To further address label uncertainty, we randomly sampled 70% of the inside-area vertices (positive) and an equal count (not percentage) of outside-area vertices from the searchlight zone (negative) to train each individual base learner (70% is based on optimization using the 50T validation dataset).

Ensembles were constructed iteratively, with each base learner trained using a different random seed on differently sampled datasets. The workflow is depicted in Fig 3 for an example area L_55b. The output of the first stage is converted to probability maps of the areas by averaging across the predictions of the 100 XGBoost classifiers for each area, and a second stage of 100 models, this time using regression decision trees (XGBoost Regressors), is trained to learn those probability maps using the same features and random vertex sampling strategy. A threshold of 0.5 is set to determine the positive and negative predictions of the regressors.

Another variant of our method was initially implemented and evaluated. Instead of using regression models (XGBoost Regressors), the same classification models (XGBoost Classifier) were used to form the ensemble in the second stage and learn from the binary predicted labels from the first stage. Although this approach inspired the final choice to use the regression model, it introduced greater deviation of the final group parcellation from the original training labels. A detailed comparison is provided in Supplementary Table S2 and Figure S6.

#### Training and evaluation

To conduct a comprehensive evaluation of the ARENA classifier, we defined two distinct models: ARENA-1 and ARENA-2, representing the trained models under the first stage and combined first and second stages, respectively. The ARENA classifiers (ARENA-1 and ARENA-2) and baseline models were trained using the training dataset (475 subjects ‘475P’; Table 1) and subsequently evaluated on multiple evaluation datasets. We assessed the performance of the areal classifier based on measures from multiple perspectives with the first perspective designed to demonstrate the classifier’s ability to accurately learn from the group parcellation (robustness), quantified using the <u>A</u>veraged <u>A</u>real <u>D</u>etection <u>R</u>atio (AADR; the same measure used in Glasser et al., 2016).

#### Dice-based reproducibility measures

Dice-based overlap is commonly used to evaluate the reproducibility of areal definitions. Accordingly, for the areal classifier, we computed the Individual <u>Di</u>ce <u>R</u>eproducibility (IDiR) across the test-retest subjects (IDiR = averaged Dice overlap across each area for one subject) and designated the median IDiR across the test-retest group as <u>T</u>est-<u>R</u>etest <u>D</u>ice <u>R</u>eproducibility (TRDR).

However, simple Dice-based overlap measures have major limitations that have not to our knowledge been critically assessed in relation to areal classifiers. If an areal classifier simply memorizes the location of each area and assigns corresponding vertices to that area in each individual, then AADR and TRDR measures would be high, but the results would add little incremental value insofar as one could apply the group labels to individuals without using an areal classifier. From our perspective, the primary objective of an areal classifier is to identify individual variability above and beyond what can be modeled with areal-feature-based cortical surface registration. To mitigate this limitation, Glasser et al. (2016) manually inspected atypical cortical topography to identify subjects observed with atypical 55b patterns in the left hemisphere. This process revealed the extent to which an areal classifier successfully captured interesting individual variability but was tedious, time-consuming and did not consider other areas. Furthermore, there is currently no evidence that all 360 areas exist in every subject—this is a topic in need of confirmation by future research. Consequently, relying on the averaged areal detection rate (AADR) can potentially lead to non-optimal methodological choices for areal classifiers.

#### Measures based on reproducible individual variability

The primary objective of the areal classifier is to capture reproducible individual variability that is not already accounted for by areal-feature-based registration. Accordingly, we introduce new metrics, the Areal Reliable Variability Score (ARVS) and individual Reliable Variability Score (iRVS), which have advantages over the AADR and TRDR measures for evaluating areal classifiers at the per-area and per-subject levels, respectively (Table 2).

**Table 2.**
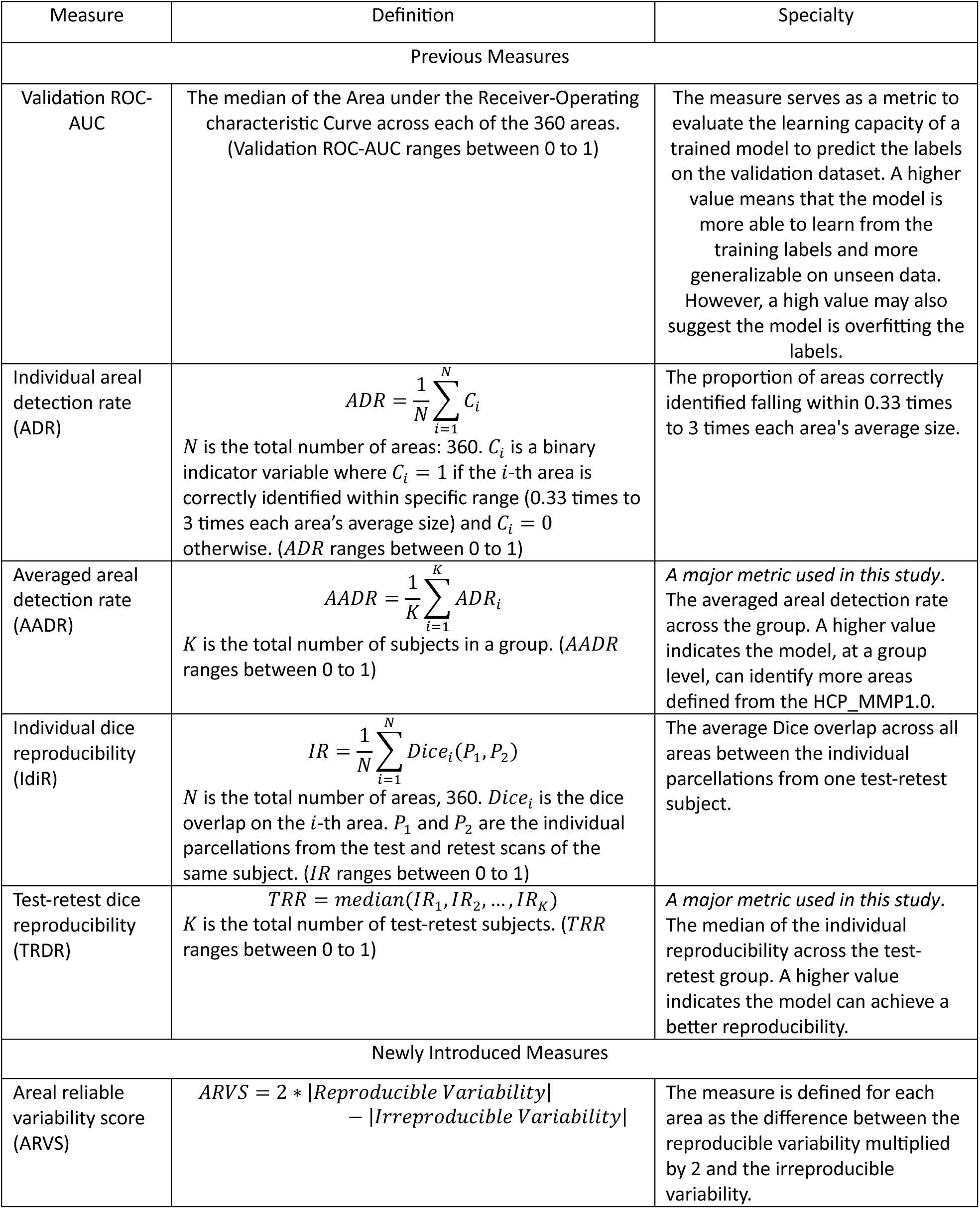
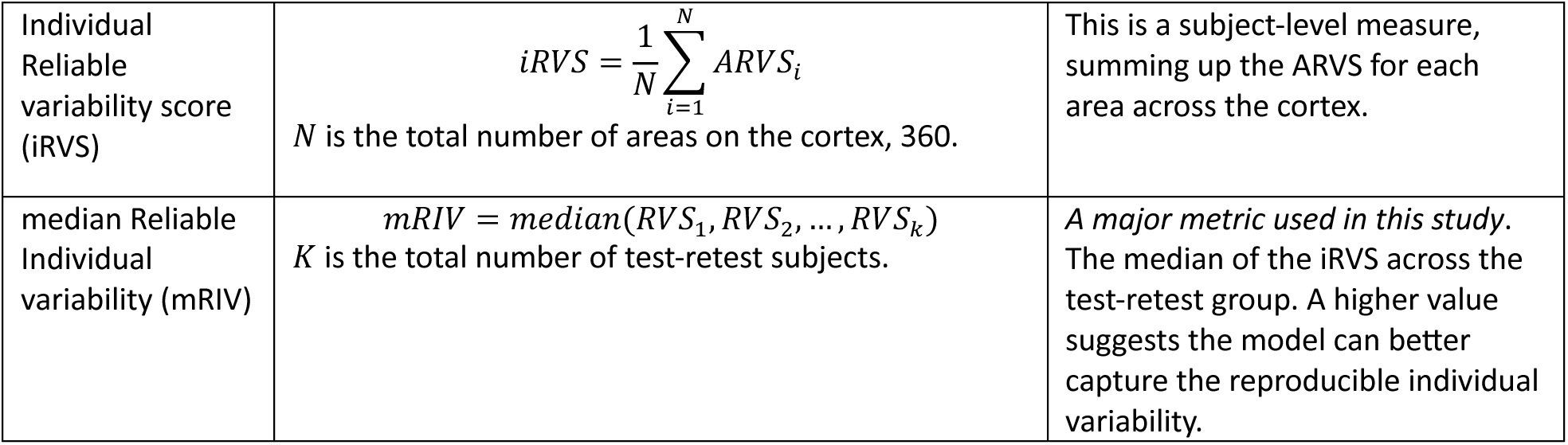
An overview of measures used to evaluate the performance of the areal classifier.

The Areal Reliable Variability Score (ARVS) is computed per area per test-retest subject, and the individual Reliable Variability Score (iRVS) is computed as the summed ARVS across the 360 areas on the whole cortex per test-retest subject. ARVS is defined for each area as the difference between the reproducible variability multiplied by 2 (because they are consensus predictions on two repeated scans) and the irreproducible variability. The reproducible variability is the count of vertices that are reproducible for a particular area in one test-retest subject (region ② in Fig 4C, which represents the reproducible variability that is predicted as inside-area by the areal classifier in both test and retest subjects and is indicated in Fig 4B by the blue and green boundaries that are overlapping, but not inside the white boundary, plus region ③ in Fig 4C, which represents the reproducible variability that is predicted as outside-area, indicated in Fig 4B by the region between the split 55b areas that is not enclosed by either a blue or green boundary, but is enclosed by the white boundary). The irreproducible variability is the count of vertices that are not reproducible (regions ④, ⑤, ⑥ and ⑦ in Fig 4C). A higher iRVS reflects a higher value of the reproducible variability and/or a lower value of the irreproducible variability, where the reproducible variability predicted by the areal classifier can be considered to be the “real” individual variability differing from the group parcellation (in this case, HCP_MMP1.0). Region ① represents the consistently reproducible predictions that match the group parcellation. To avoid evaluation bias introduced by the group parcellation, this region is considered neutral, neither helping nor hurting a classifier’s performance.

**Figure 4.**
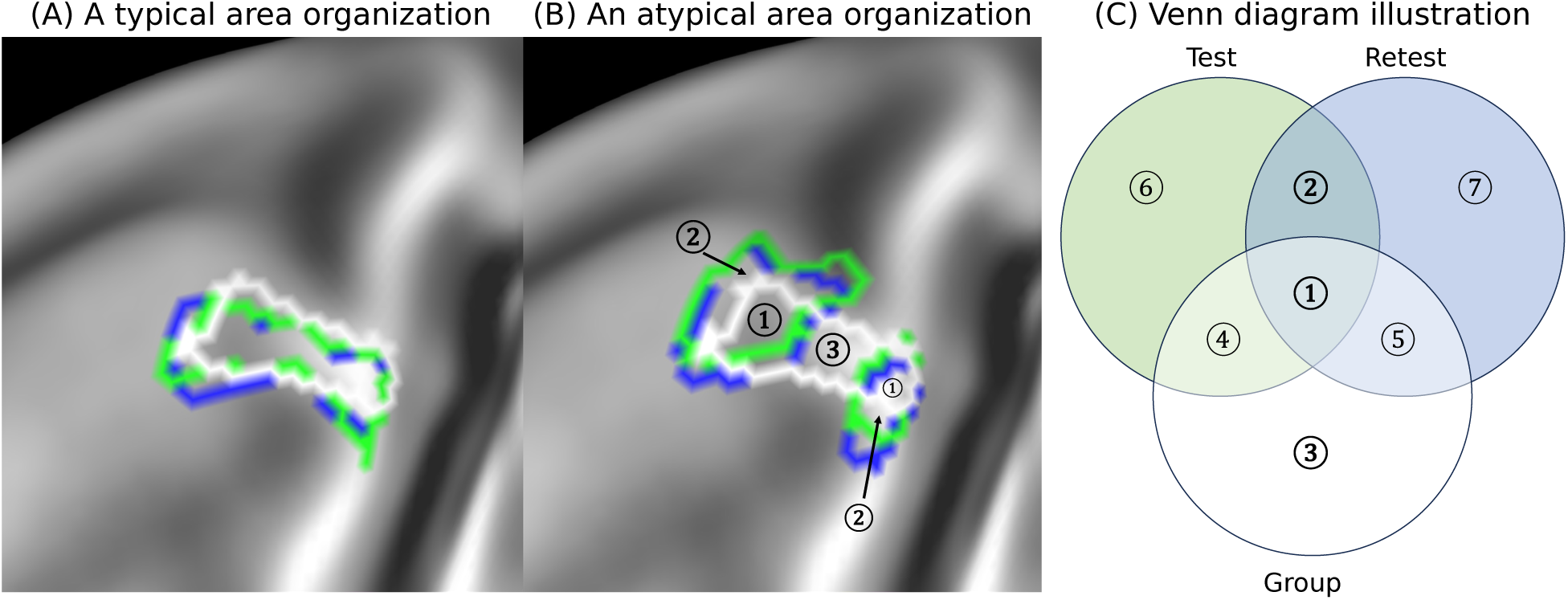
An exemplar to demonstrate the measures leading to an overall performance evaluation. The predicted L_55b from test (green) and retest (blue) scans and the label from HCP_MMP1.0 (white) are shown for (A) a subject (111312) with typical organization and (B) a different subject (149337) with atypical organization. The three parcellations for one area can be framed by a Venn diagram (C) as three sets, each of which contains the vertices within the parcellation (parcels or predicted areas), where region ① stands for the reproducible training labels (present in all 3), regions ② and ③ stand for the reproducible variability predicted by the areal classifier, while the rest of the regions stand for irreproducible variability that should be minimized. These three main regions ①, ② and ③ are marked in panel (B) for corresponding information. The Dice overlap in (A) for the area predicted in the test-retest subject is 0.9 (Dice between blue and green regions) and the areal detection indicator for this area is 1 (because in both test-retest predictions, the predicted area size falls between 0.33x and 3x of the group averaged size). The Dice overlap in (B) for the area predicted in the test-retest subject is 0.9 and the areal detection indicator for this area is 1. To compute the reproducibility for this test-retest subject, the Dice overlap for each area is averaged across all the areas. To compute the areal detection rate for each subject, the areal detection indicator is averaged across all the areas. The individual Reliable Variability Score (iRVS), which is sensitive and has a great dynamic range, is computed as 2 ∗ (|②| + |③|) − (|➃| + |⑤| + |⑥| + |⑦|). For the exemplar subjects, ARVS=49 for case A and ARVS=109 for case B.

We consider iRVS as the best overall measure for evaluating different areal classifiers. To reveal the overall effect in the test-retest group, the median of iRVS across the test-retest group is reported as <u>m</u>edian <u>R</u>eliable Individual <u>V</u>ariability (mRIV).

Table 2 describes the various evaluation metrics used in this study.

#### Implementation

The XGBoost learner was implemented using the XGBoost module with scikit-learn wrapper (Chen & Guestrin, 2016) and was trained on CPU only. The individual parcellation pipeline is publicly available as part of the HCP Pipelines https://github.com/Washington-University/HCPpipelines.

## 3. Results

The Results are divided into two major sections, each with multiple subsections. Sections 3.1 - 3.4 demonstrate the methodological improvements achieved by our ARENA classifier, which begins with the effectiveness of ensemble learning in the first stage of the training schema (#3.1; ARENA-1), highlighting the benefits of ensembling and random subsampling for achieving robust, reproducible and reliable individual parcellations. Next, we investigate the impact of different types of RSN decomposition on individual parcellation performance, examining the relationship between RSN decomposition and accuracy, reproducibility, and the ability to capture reliable individual variabilities (#3.2; ARENA-1). Section 3.3 directly compares the ARENA classifier with the original MLP model (ARENA-1 and ARENA-2). Section 3.4 demonstrates the generalizability of the ensemble classifier when applied to unseen task fMRI data (ARENA-1 and ARENA-2).

The second section includes two sets of important neurobiological findings: 1) the HCP_MMP1.0_1071_MPM refined group parcellation (#3.5), using an advanced areal classifier and larger sample size to improve multimodal group parcellation along with examples (#3.6) to demonstrate individual variability in test-retest subjects and 2) a summary of atypical 55b organization in both hemispheres (#3.7), revealing two novel types of topological organization and contributing to a summary of 55b typical and atypical organization for the HCP-YA 1071 subjects.

### 3.1 Effect of ensemble learning

To demonstrate the effectiveness of ensemble learning in enhancing the performance of the ARENA classifier, we trained five ARENA-1 models using the same dataset, consisting of 475 subjects (475P) with spatial ICA feature maps (n=48 for d=76 WD5) as RSN maps. The models considered were: 1) A single XGBoost model trained on subjects that are among the most similar 75% compared with the group average feature vector (Column #1 in each subplot of Fig 5), 2) An ensemble of 20 XGBoost models trained on the best matching 75% of subjects (Column #2), 3) An ensemble of 20 XGBoost models trained on all subjects (Column #3), 4) An ensemble of 100 XGBoost models trained on the best matching 75% of subjects (Column #4), and 5) An ensemble of 100 XGBoost models trained on all subjects (ARENA-1; Column #5). We evaluated the five models based on four metrics: a) Validation ROC-AUC, to assess the model’s learning capacity (Fig 5A), b) average areal detection rate (AADR) on the 475V evaluation dataset (Fig 5B blue) to evaluate the classifier’s performance on an unrelated group of subjects, AADR on the 475P training dataset (Fig 5B orange) to evaluate the classifier’s learning capability, c) test-retest Dice reproducibility (Fig 5C), and d) reliable individual variability, expressed as the median Reproducible Individual Variability (mRIV). Strikingly, the ARENA-1 with random subsampling (Columns #2) performed substantially better than the single XGBoost (Column #1) trained on the same top 75% group in all four measures. When more subjects are involved in the training process, including less well-matched subjects (Column #3), the model slightly outperformed the version applied to only the well-matched subjects with the same number of base learners (Column #2) in all four metrics. The performance difference was more substantial for the reliable individual variability on the test-retest 45 group. With more base learners and only best matching subjects (Column #4 vs. Column #2), a small performance reduction is seen in mRIV with virtually no change in other metrics. A combined treatment (ARENA-1; Column #5 in orange) with more base learners and all subjects involved in training demonstrated a slight overall performance gain in all the four metrics relative to fewer learners (Column #3). In summary, treating areal classification as a weakly supervised learning problem, particularly when employing ensemble learning and random subsampling, is a promising approach.

**Figure 5.**
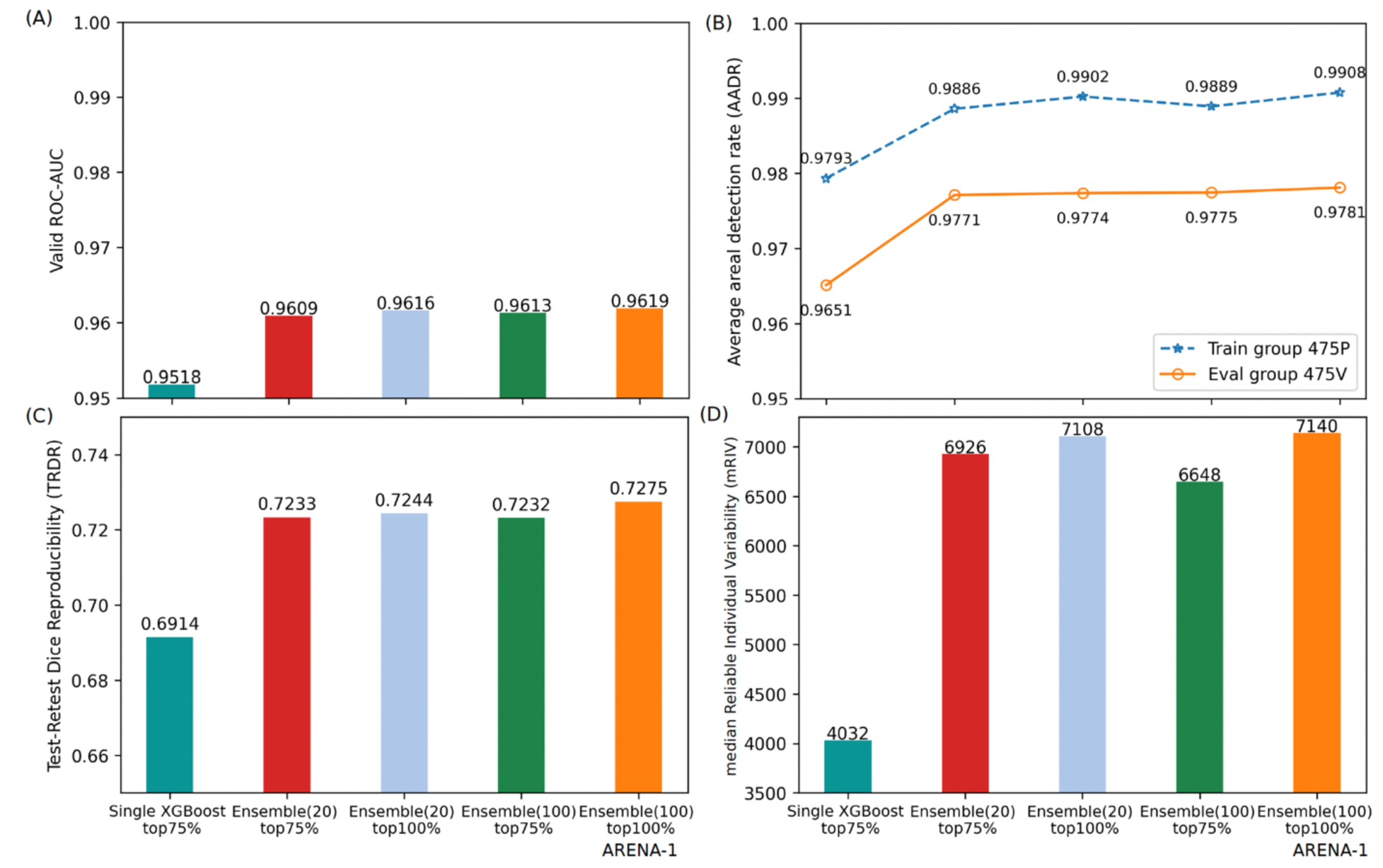
Evaluation comparison on ensemble learning (ARENA-1). (A) Validation ROC-AUC. (B) Averaged Areal Detection Rate (AADR) on the 475V evaluation dataset (orange) with unrelated subjects to the 475P training group (blue). (C) The Test-Retest Dice Reproducibility. (D) The median Reliable Individual Variability (mRIV – see Table 3).

**Table 3.** Performance evaluation under four RSN decomposition types: sICA, tICA, PFM-DR and PFM-WR. The best two methods are shown in bold.

| RSNs | models | median Reliable Individual variability | Averaged areal detection rate on 475V (%) | Test-Retest Dice Reproducibility (%) |
| --- | --- | --- | --- | --- |
| sICA d137 from Glasser et al, 2016 | MLP | 3048 | 95.28 | 68.5 |
| sICA d137 from Glasser et al., 2016 retrained | MLP | 3128 | 97.23 | 71.6 |
| sICA d76 | MLP | 5238 | 96.95 | 70.4 |
|  | ARENA-1 | 7140 | 97.81 | 72.8 |
|  | ARENA-2 | 7614 | 97.44 | 71.4 |
| tICA d76 | ARENA-1 | 7276 | 97.91 | 73.3 |
|  | ARENA-2 | 7990 | 97.60 | 72.0 |
| PFM-DR d76 | ARENA-1 | 8876 | 97.54 | 72.6 |
|  | **ARENA-2** | **9786** | 97.28 | 71.5 |
| **PFM-WR d76** | ARENA-1 | 7416 | 98.10 | 73.8 |
|  | **ARENA-2** | **8380** | 97.81 | 73.3 |
| sICA d92 | ARENA-1 | 5632 | 98.01 | 73.6 |
|  | ARENA-2 | 6784 | 97.65 | 72.0 |
| tICA d92 | ARENA-1 | 6060 | 98.13 | 74.1 |
|  | ARENA-2 | 7520 | 97.78 | 73.0 |
| PFM-DR d92 | ARENA-1 | 8114 | 97.76 | 72.7 |
|  | ARENA-2 | 9162 | 97.45 | 72.0 |
| PFM-WR d92 | ARENA-1 | 5928 | 98.32 | 75.0 |
|  | ARENA-2 | 7450 | 98.04 | 74.2 |

### 3.2 Effect of different types of RSN maps

RSN (Resting-State Network) maps contain essential functional information and serve as a major determinant of areal boundaries, albeit to varying degrees for different areas (see Introduction). To compare various RSN candidates, we trained ARENA-1 (not ARENA-2, for the purpose of RSN comparison only) with 100 base learners with all subjects on the feature maps (the best approach from the preceding section), where the RSN maps were based on five types of representation: sICA (Columns #1 & #2 in Fig 6), tICA (Columns #3 & #4), PFM-Dual Regression (PFM-DR; Columns #5 & #6), PFM-Weighted Regression (Column #7), and PFM origmap (Columns #8 & #9), while keeping the remaining features unchanged. Two different dimensionalities of feature maps were considered: 92 and 76, derived from group spatial ICA after fitting 5 and 6 Wishart Distributions, respectively, to represent RSNs more broadly while considering varying numbers of components for ICA/modes for PFM. Four Wishart Distributions, resulting in 190 components, was visually inspected and found to have substantial component over-splitting and worse performance in identifying reproducible individual variability (mRIV of 3376), suggesting that it was allowing overfitting in the classifier. Furthermore, smoothing the tICA d=92 maps by 4 mm FWHM led to less reliable individual variability (mRIV of 5790) compared with the unsmoothed version (tICA d92 WD5 in panel D from Fig 6; mRIV of 6060).

**Figure 6.**
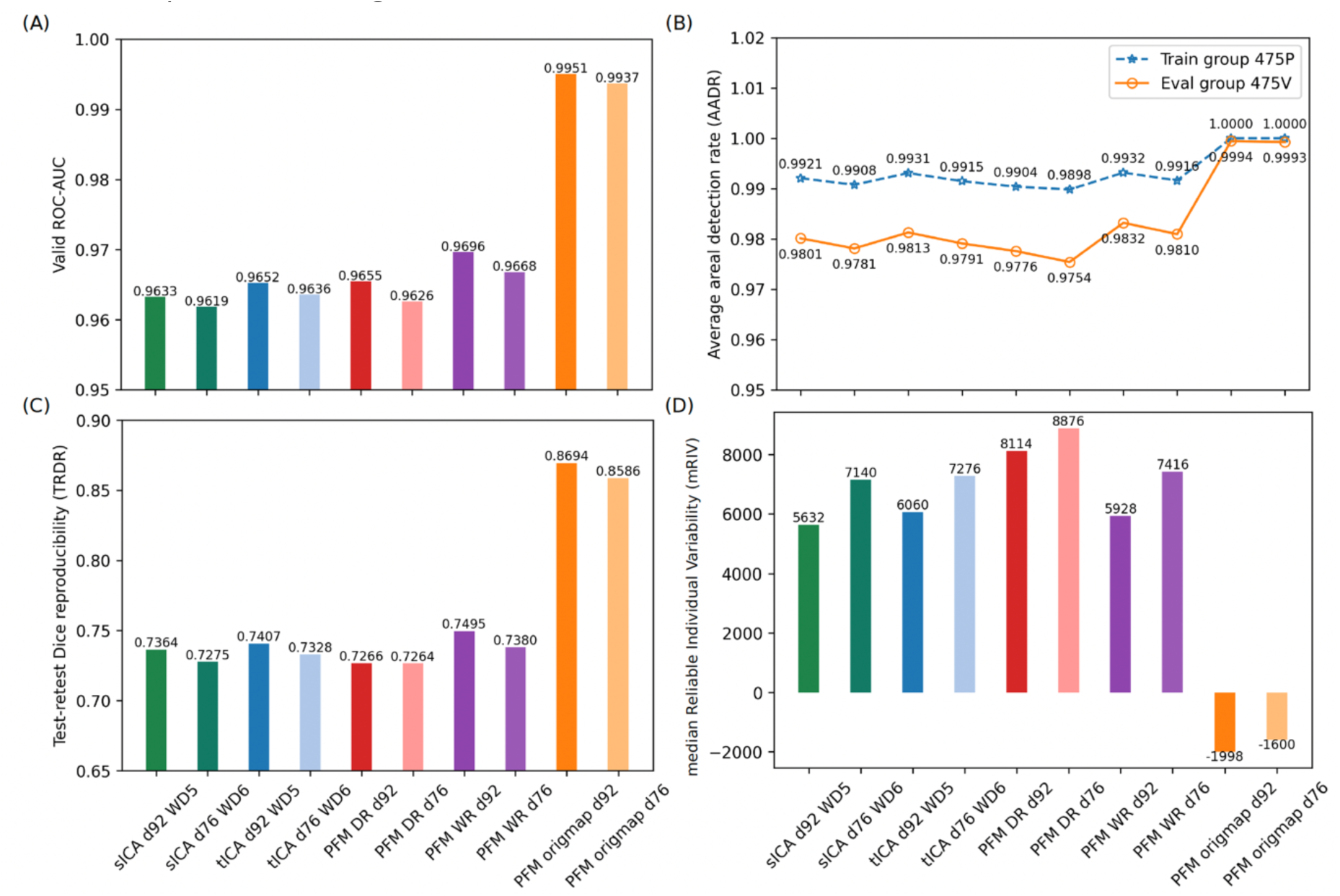
Evaluation comparison on different types of RSN maps: sICA (green), tICA (blue), PFM-Dual regression (PFM-DR; red/pink), PFM-Weighted Regression (PFM-WR; purple), and PFM origmaps under two dimensionalities: 92 and 76, estimated from WF5 and WF6 on group sICA from HCP-YA 1071 3T subjects (orange). The four metrics consisted of (A) Validation ROC-AUC on the 50T validation dataset, (B) Averaged areal detection rate (AAPR) on the 475P training and 475V evaluation datasets, (C) The Test-Retest Dice Reproducibility (TRDR), and (D) median Reliable Individual Variability (mRIV).

In comparing results for sICA and tICA maps, tICA maps demonstrated superior performance in all the four measures: validation ROC-AUC, average areal detection rate (AADR), test-retest reproducibility (TRR), and reliable individual variability. The overall difference between sICA and tICA may be attributed to the higher Signal-to-Noise Ratio (SNR) of tICA features, resulting in a better fit to the training labels and a greater ability to capture reliable individual variability. As both sICA and tICA maps were generated using a two-step process, group-level decomposition and regression into individual dense timeseries, these maps were more comparable to PFM-WR than to the other PFM-related maps. The enhanced performance of tICA suggests that features containing all functional connectivity information produce a more robust and accurate individual parcellation than sICA-based features, where some of such information is present in correlations between timeseries (i.e., the network matrix) and is not seen by the areal classifier.

Using a Bayesian framework to derive RSN maps, PFM origmaps exhibited substantial improvements in validation ROC-AUC, averaged areal detection rate and test-retest reproducibility. However, this approach had the worst reliable individual variability (mRIV; panel D from Fig 6), primarily due to features that led the classifier to overfit to the training labels (i.e., the group parcellation). This pattern arose because PFM origmaps contained high-contrast features, causing the classifier to learn incorrect features for certain areas (e.g., L_55b) as illustrated in Fig S1-3. After feature selection and subject filtering based on feature similarity, the model trained on PFM origmaps could capture individual variability (e.g., a split 55b), but performed significantly worse in averaged areal detection rate and test-retest reproducibility (Refer to Table S1 and Fig S5). The poor performance of PFM origmaps in capturing reliable individual variability suggested that it should not be used as the RSN features in areal classification.

One variant of PFMs, PFM-DR, consistently demonstrated superior performance in reliable individual variability compared to sICA and tICA, showing that it was the best RSN decomposition for capturing reproducible individual variability. However, it was not chosen as the optimal RSN decomposition because generalizing to unseen subjects or types of fMRI data requires rerunning the entire group+individuals’ decomposition whenever new subjects are added.

On the other hand, the workaround solution, PFM-WR, led to a worse performance in reliable individual variability compared to PFM-DR, but outperformed both sICA and tICA in reliable individual variability, validation ROC-AUC, averaged areal detection rate (AADR), and test-retest reproducibility (TRDR). These findings show that PFM decompositions, which do not enforce spatial or temporal independence, are superior to sICA and tICA as features, which do enforce spatial or temporal independence, for cortical areal classification, even when not optimally derived using a full run of PROFUMO.

Additionally, the decrease in dimensionality (from 92 to 76) for sICA, tICA, PFM-DR, PFM-WR, and PFM-origmaps RSN decomposition was associated with increased reliable individual variability and reduced validation ROC-AUC, average areal detection rate and test-retest reproducibility because the reduced complexity of RSN features makes it harder for the model to memorize the group parcellation. As capturing reliable individual variability is the main purpose of the areal classifier, d=76 is preferred over d=92 (also WD6 was the final choice for the official release of HCP-YA 3.0 and HCP-Lifespan 3.0), resulting in the top two candidates for RSN decompositions being PFM-DR d=76 and PFM-WR d=76.

### 3.3 Benchmark comparisons

We carried out a benchmark comparison to assess the performances of the ARENA-1 and ARENA-2 classifiers relative to the previously published (baseline) MLP model with sICA at d=137 and updated best sICA dimensionality of d=76. This comparison was conducted for ARENA-1 and ARENA-2 for four RSN decomposition types under d=76 and d=92: 1) sICA RSN maps, which were used for the original MLP model, 2) tICA RSN maps, plus 3) PFM-DR maps and4) PFM-WR maps - the top two candidates from the previous section. The first comparison between the MLP and ARENA classifiers provides fundamental evidence of how weakly supervised learning enabled by the ARENA classifier benefits areal classification. The second comparison across RSN types and the two ARENA versions highlights the optimal approach for areal classification.

Evaluations were based on three measures shown in Table 3: 1) Reliable Individual Variability Score (Table 3), 2) Averaged areal detection rate, and 3) Test-retest Dice Reproducibility. The first and the third measures were computed on the 45 test-retest subjects, whereas the second was computed on the 475V evaluation group.

Table 3 shows that retraining the MLP model with the original d=137 features resulted in modest improvement, mostly in detection rate and dice overlap, while the MLP trained on d=76 substantially improved the mRIV score with little sacrifice to detection rate or dice, suggesting less overfitting to the group labels. Section sICA d76 shows that, for the sICA RSN decomposition, the ARENA-1 model outperformed the MLP model on all metrics, suggesting significant improvement resulting from the model architecture involving weakly supervised learning using ensemble and random sampling. When comparing the RSN decomposition methods using the ARENA-1, sICA and tICA (d=92 and d=76), tICA outperformed sICA in every metric, expanding on findings from the previous section. PFM-DR outperformed tICA on score-based Reliable Individual Variability, but neither Average Areal Detection Rate nor Test-Retest Dice Reproducibility. Additionally, the lower dimensionality d=76 achieved better reproducible individual variability metrics than d=92 across all the types of RSN decomposition. The best two ARENA-1 models are PFM-WR d=76 and PFM-DR d=76, which is also true for ARENA-2. ARENA-2 shows better reproducible individual variability than ARENA-1, illustrating the effectiveness of incorporating a second round of training to address label uncertainty, which utilizes the pseudo-target generated from the ARENA-1 model. Although PFM-DR with ARENA-2 and d=76 shows the highest performance, because it is difficult to extend to new individuals and groups, we chose PFM-WR as the approach with the current greatest utility.

We conducted an additional set of benchmark comparisons to demonstrate the overall methodological improvement using spatial maps of areal detection rates, areal probability ratios, and areal reliable variability ratios (Fig 7). In this comparison, MLP with sICA d=137 (rows 1, 4, 7 in Fig 7) was compared with ARENA-2 with PFM-DR d=76 (rows 2, 5, 8) and PFM-WR d=76 (rows 3, 6, 9) all using the same 475V evaluation (rows 1-6) and 45 test-retest subjects (rows 7-9). For the areal detection rate (rows 1, 2, 3), ARENA-2 exhibited better overall performance, with rows #2 and #3 having fewer red (low detection) areas, mainly in anterior cingulate cortex. The areal probability ratio (rows 4-6) describes the fraction of the areal probabilities captured by the maximum probability map for each area (with 1 being a probability map having no spread outside of the MPM). The maps are visually consistent across models, suggesting that all three areal classifiers are able to make similar probabilistic maps (and generally agree on the amount of individual variability of most areas). The areal reliable variability score (rows 7 – 9) was computed for each area for every model, where a higher value indicated a greater ability to capture reliable individual variation from the group parcellation.

**Figure 7.**
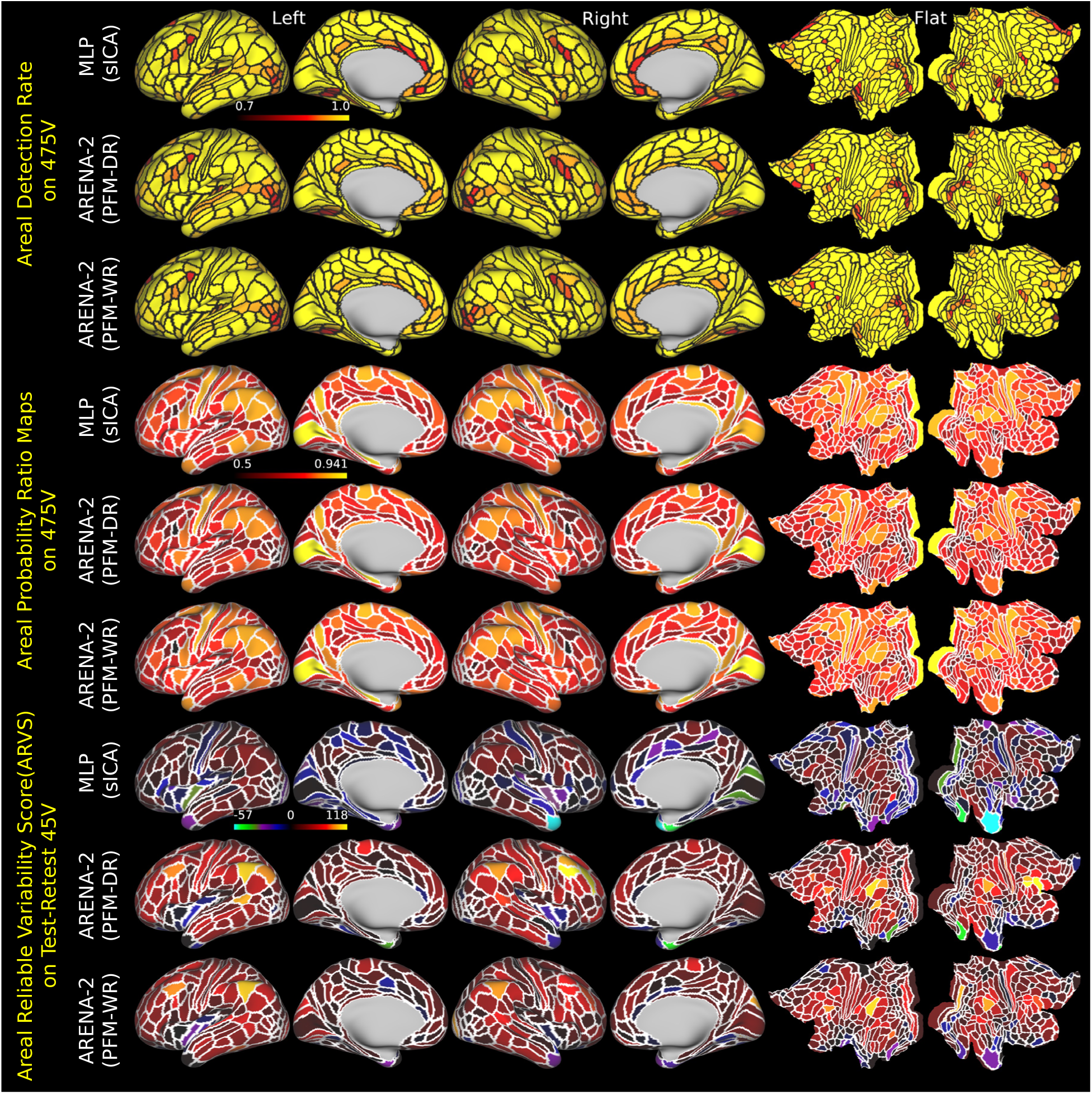
Areal detection rates, Areal probability ratio and Areal reliable variability ratio. Each section comprises three rows, corresponding to the results from the baseline MLP (sICA d=137), ARENA-2 (PFM-DR) and ARENA-2 (PFM-WR) runs with d=76. Areal Detection Rate (Rows #1 to #3): Individual areal detection rates are displayed for the evaluation group of 475 subjects. Detection rates are scaled from 0.7 to 1.0, with minimum rates of 0.75 (Row #1), 0.68 (Row #2), 0.78 (Row #3). The averaged areal detection rates are displayed in Table 3. Areal probability ratio (Rows #4 to #6): the proportion of each area’s probability map inside a given method’s group MPM divided by the sum of the probabilities both inside and outside that method’s group MPM. The maps are scaled from 0.5 to 0.9407 (the maximum value across the three datasets). Areal reliable variability ratio (Rows #7 to #9): the median of the areal reliable variability score scaled from −57 to 118 (the minimum and maximum value across the four maps) are illustrated for the 45 test-retest subjects, with the values for L_55b of 19 (Row #7), 46 (Row #8), and 40 (Row #9). Values less than 0 indicate areas for which the areal classifier’s result is less reliable than simply imposing the group parcellation (because true topological individual variability in these areas is low, and they are already well aligned across subjects by areal feature-based registration).

The areal probability ratio and areal reliable variability score (ARVS) show opposite patterns because they are measuring similarity and dissimilarity compared to the group-level parcellation. For instance, V1 areas are less variable across subjects, leading to a high areal probability ratio (the maximum yellow; meaning that V1 areas are closely aligned for each individual in the 475V group) and a low areal reliable variability score (darker). The ARENA-2 together with PFM RSN maps (with more yellow areas in rows 8 and 9) capture reliable individual variability better than the MLP with sICA maps. Notably, PFM-DR captures more reliable individual variability than PFM-WR, while PFM-WR outperforms sICA. When including practical considerations, we consider PFM-WR to currently be the most useful RSN decomposition method, as it combines the strengths of both PFM and ICA-based methods: with the greatest abilities to capture reliable individualized patterns as well as to generalize to unseen subjects or new fMRI runs.

### 3.4 Generalizability to a different fMRI paradigm

We assessed the generalizability of the ARENA models (1 & 2) on task fMRI data using three RSN decomposition types: sICA, tICA, and PFM-WR (PFM-DR was excluded based on the need to retrain the classifier, creating practical issues). Even though group ICA and PFM are typically applied to resting-state fMRI data, we can generate individualized spatial maps for arbitrarily concatenated or single task fMRI runs based on the resting-state group decompositions in both ICA and PFM. Our goal was to validate the model’s generalizability by using task fMRI data that the model had not encountered during the training process. Using d=76 resting state decompositions, we trained the MLP (red in Fig 8) using sICA, ARENA-1 model (teal, yellow and green bars in Fig 8) using sICA, tICA, and PFM-WR, and ARENA-2 model using tICA (blue bar in Fig 8) and PFM-WR (purple bar in Fig 8) on the 475P training dataset. We then evaluated these trained models on two evaluation datasets, 475 subjects (475V) unrelated to the 475P training subjects and 45 test-retest subjects using both resting-state fMRI (left six bars in each panel from Fig 8), and concatenated task fMRI (right six bars in each panel).

**Figure 8.**
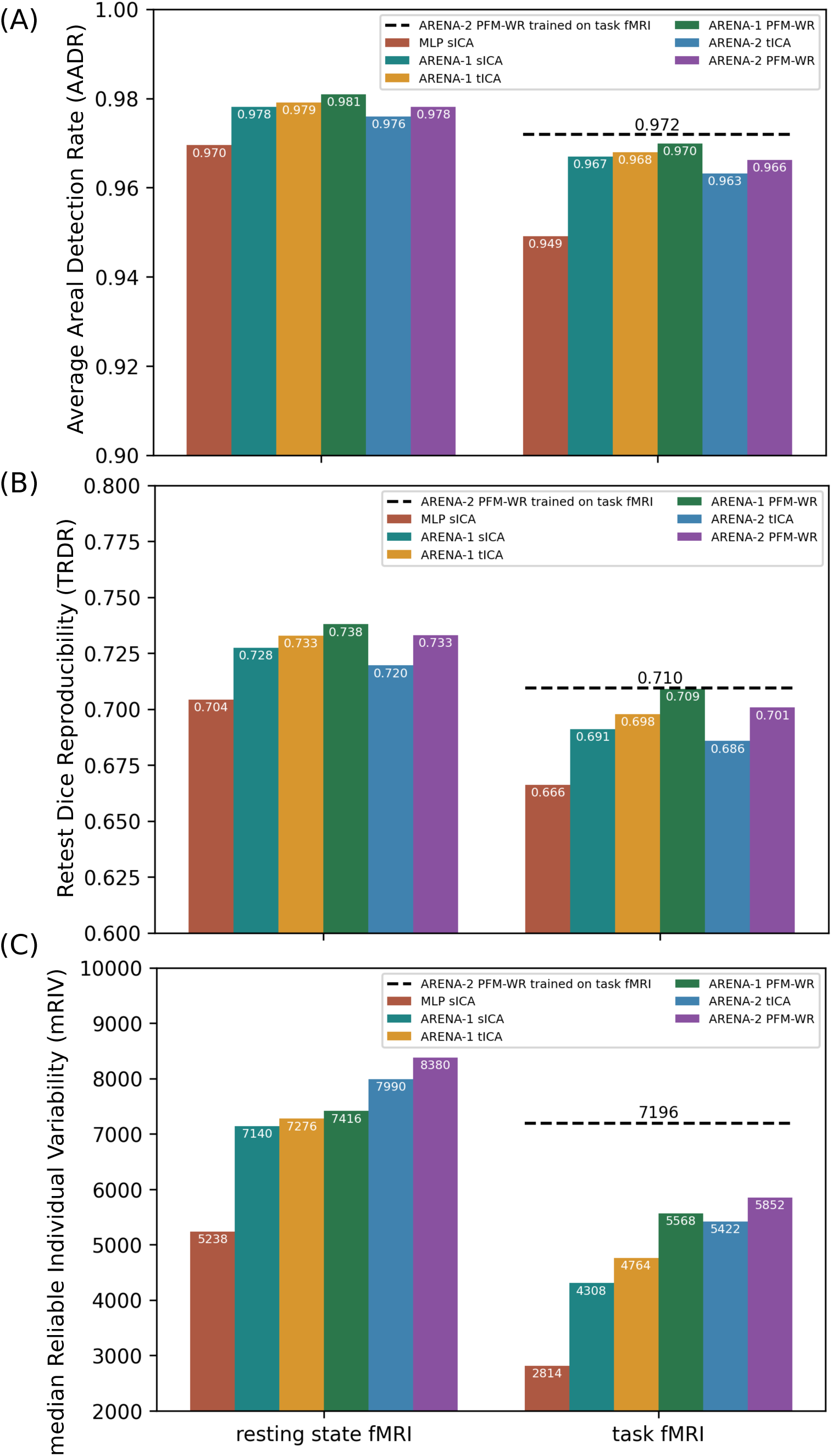
Generalizability of d=76 resting state trained models to task fMRI data. Averaged areal detection rate on 475V group (A). Dice reproducibility on test-retest individual parcellations (B). The score-based Reliable Individual Variability (mRIV). The best estimate of the upper bound for the task fMRI data is shown as a dashed black line for each panel using an ARENA-2 model trained on PFM-WR features derived from the task (at the task fMRI’s natural WD6 dimensionality of d=65).

We computed the averaged areal detection rates for the 475V group (Fig 8A), test-retest reproducibility (Fig 8B), and score-based reliable individual variability (Fig 8C). Consistently, the ARENA models outperformed the MLP using sICA in all three measures when comparing across the same RSN decompositions, demonstrating superior generalizability.

In summary, the ARENA models with PFM-WR RSN maps were most generalizable to task fMRI not involved in the training process. An empirical upper bound of each measure for task fMRI is provided in each panel obtained from a retrained ARENA-2 model using PFM-WR-based RSN features derived from the task fMRI (d=65, the WD6 optimum dimensionality for the task data). Comparing the best estimations from the task and resting state fMRI derived RSN features (the dashed black line and the purple bar), the task fMRI is less informative at capturing reproducible individual variability with worse averaged areal detection rate and test-retest reproducibility.

ARENA-2 obtained the better reliable individual variability (the blue and purple bars In Fig 8C), consistent with its better performance in the resting-state fMRI.

### 3.5 HCP-YA Multimodal Parcellation Maximum Probability Map (MPM) on 1071 3T subjects

Using the more advanced areal classifier and larger HCP-YA study group, a refined multimodal parcellation on the cortex was generated, named HCP_MMP1.0_1071_MPM. We trained the ARENA-2 with 100 base learners using the PFM-WR RSN d=76 and resting-state fMRI on the 475P training group. The trained model was then applied to the entire set of 1071 subjects and their individual parcellations were combined to create the HCP_MMP1.0_1071_MPM group parcellation using the group maximum probability map (MPM; Fig 9 row #1, row #3 in red). We also obtained the MPM group parcellation with motor/sensory subregions (Fig 9 row #2 in black) and subarea group parcellation (row #3 in green) from the intersection between the HCP_MMP1.0_1071_MPM (row #1) and subregion parcellation (row #2). As originally described in Glasser et al., 2016 Supplementary Neuroanatomical Results section 6 and Figures 7 and 8, the sensori-motor cortex (cortical areas 4, 3a, 3b, 1, and 2) can also be subdivided into sensorimotor subregions for the face/head, eye, upper extremity, trunk, and lower extremity. The intersection between the cortical areas and somatotopic subregions produces a set of somatotopic sub areas of each cortical area. We have classified the subregions as well as the cortical areas, enabling their intersection to be used to produce subareas in any individual. The Dice overlap between the original semi-automated parcellation (HCP_MMP1.0_210P_Orig, also the training label for the areal classifier; row #3 in blue) and the 1071 group MPM was 0.888, indicating that modest adjustments were needed to better fit the data, similar to the comparison between the 210P group MPM and HCP_MMP1.0_210P_Orig (Glasser et al., 2016). Row #4 provides a visual comparison between the borders, with purple representing overlapping vertices, once again demonstrating a strong correlation. To evaluate area size change, we calculated the log_2_ of the ratio between each area’s size in HCP_MMP1.0_1071_MPM and the original size in HCP_MMP1.0_210P_Orig, 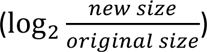.

**Figure 9.**
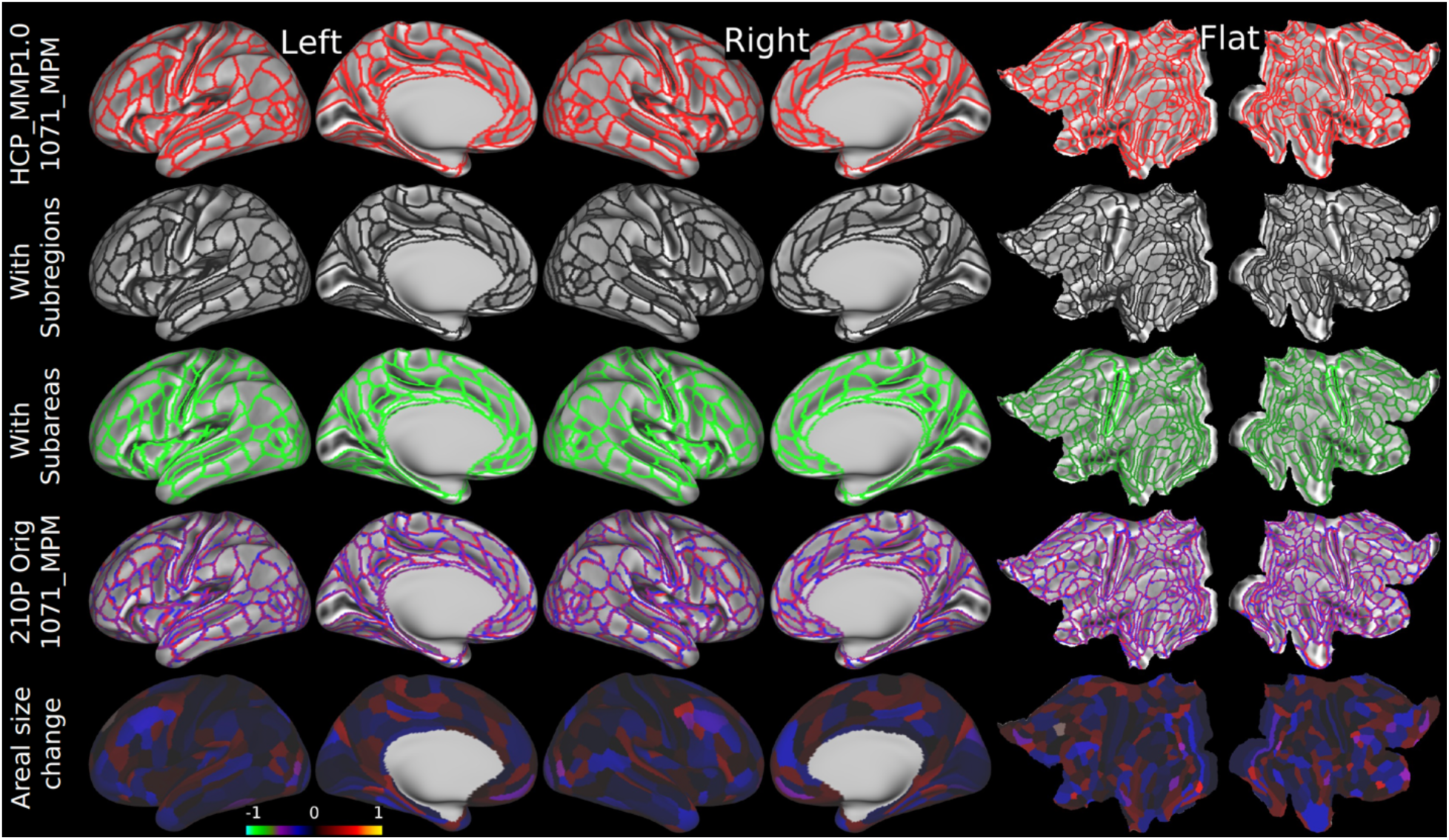
The HCP_MMP1.0_1071_MPM group parcellation, group parcellation with subregions, group parcellation with subareas, parcellation reproducibility, and areal size bias compared with the HCP_MMP1.0_210P_Orig version are shown. Row #1 shows HCP_MMP1.0_1071_MPM, which was generated using the ARENA-2 model trained on the 475P dataset with the PFM-WR WF6 RSN decomposition and applied to the HCP-YA 1071 3T subjects (HCP_MMP1.0_1071_MPM_CA for <u>C</u>ortical <u>A</u>reas). Row #2 shows the MPM parcellation with subregions (in black; HCP_MMP1.0_1071_MPM_SR for <u>S</u>ub-<u>R</u>egions) and row #3 shows the group parcellation with subareas (the intersection between row #1 and #2; in green; HCP_MMP1.0_1071_MPM_SA for <u>S</u>ub-<u>A</u>reas). Row #4 represents the parcellation reproducibility between HCP_MMP1.0_1071_MPM (in red) and HCP_MMP1.0_210P_Orig (in blue). Overlaps are displayed in purple. Row #5 displays the log2 ratio between the area size in HCP_MMP1.0_1071_MPM and that in HCP_MMP1.0_210P_Orig. A value of 1 indicates that the area in HCP_MMP1.0_1071_MPM is twice the size of its counterpart in HCP_MMP1.0_210P_Orig. A ratio of 0 signifies that the sizes are equivalent, and a ratio of −1 means that the area is half the size.

A value of −1 indicates that the HCP_MMP1.0_1071_MPM is half of the original size, whereas a value of 1 indicates a size twice that of the original. The areas with the greatest positive size change (larger sizes) were L_47m_ROI (0.37) and R_VMV1_ROI (0.45) in the left and right hemispheres, respectively. Conversely, the areas with the greatest negative size change (smaller sizes) were L_L01_ROI (−0.30) and R_9p_ROI (−0.33) in the left and right hemispheres. Notably, no area was missing from HCP_MMP1.0_1071_MPM, indicating a reliable replication of the group parcellation HCP_MMP1.0_210P_Orig.

### 3.6. Examples of Individual variability in test-retest subjects

Fig 10 illustrates some of the diversity in both reproducible and irreproducible individual variability using 10 exemplar cortical areas in each of three HCP test-retest subjects processed using the PFM-WR d76 version of the ARENA-2 classifier. The top row shows the group average parcellation (Fig 10A) plus parcellations for test (Fig 10B) and retest data (Fig 10C) in subject 111312. Inspection of corresponding areas on these maps reveals many differences between the group average and the individual test and retest area boundaries. To facilitate systematic comparisons, Fig 10D shows areal boundaries for the group average (black contours) and the 111312 test and retest data (yellow and red contours, respectively) for 10 cortical areas visible on lateral inflated left hemisphere views. Figs. 10E and F show test and retest areal boundaries for the same 10 areas in exemplar subjects 115320 and 103818 overlaid on group average parcel outlines in black. Places where yellow and red contours deviate consistently from the blue contour are indicative of reproducible individual variability (analogous to where blue and green test-retest contours jointly deviate from white group average contours in Fig. 4), whereas differing trajectories of yellow and red contours indicate irreproducible individual variability.

**Figure 10.**
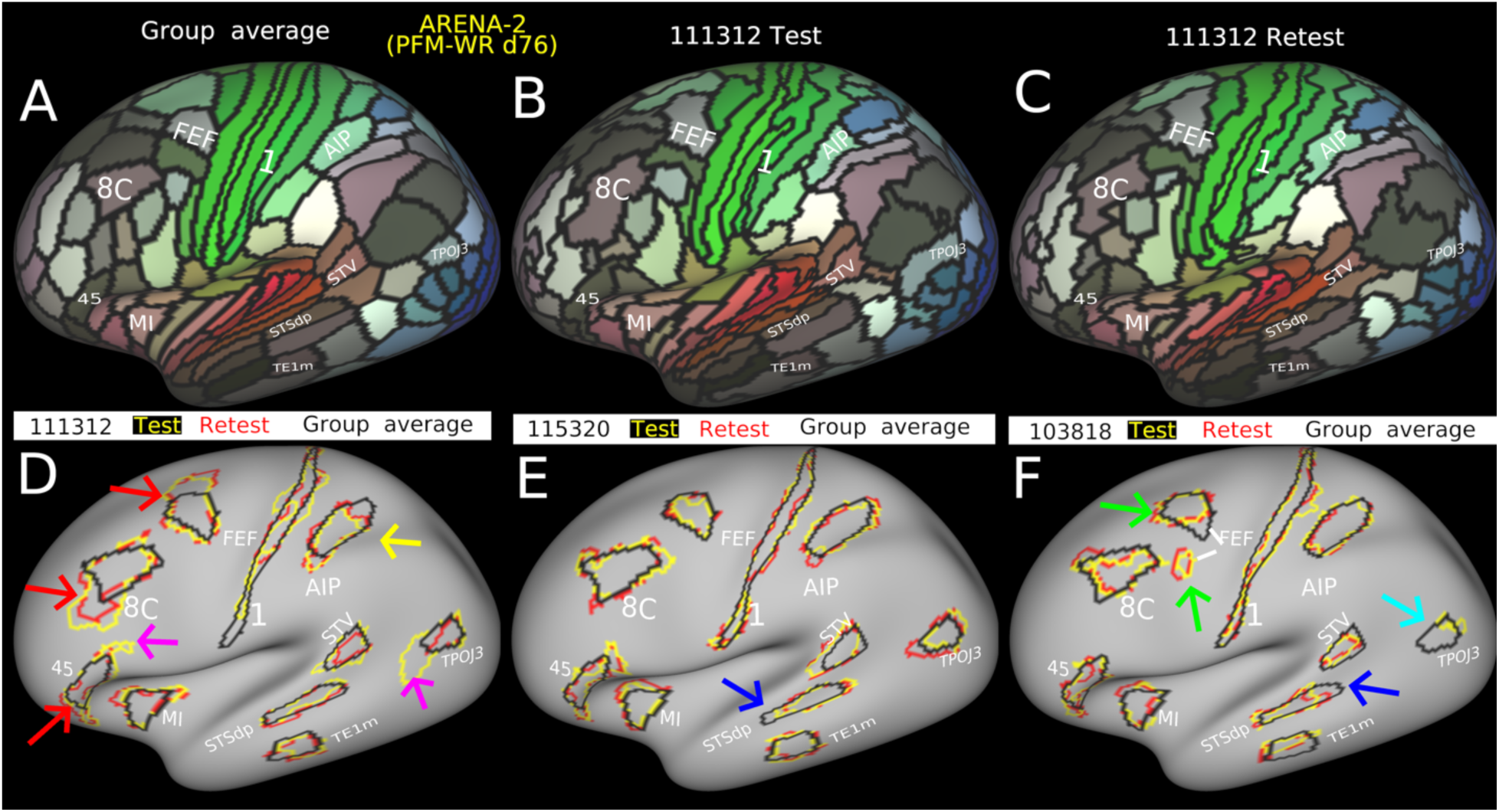
Exemplar maps of individual variability using the ARENA-2 PFM-WR d76 areal classifier. (A) Group average of HCP 1071 subjects with 10 exemplar areas labeled that are visible on a lateral view of the inflated left hemisphere surface. (B) and (C) Parcellations of subject 111312 test and re-test datasets. (D - F) Areal boundaries for the group average (blue contours) and individual-subject test and retest datasets (yellow and red contours, respectively) for 10 cortical areas from subjects 111312 (D), 115320 (E), and 103818 (F).

Examples of strong test-retest reproducible variability include ventral extensions of areas 10C and 45 and a medial extension of FEF in subject 111312 (red arrows, Fig 10D), a reproducible split representation of FEF in subject 103818 (green arrow, Fig 10F), a reproducibly smaller extent of STSdp in subjects 115320 and 103818 (blue arrows, Fig 10E &F), a reproducibly shifted extent of AIP in subject 111312 (yellow arrow, Fig 10D), and a TPOJ3 that is much smaller than the group average in the retest data for subject 103818 and missing altogether in the test data (aqua arrow in Fig 10F). Examples of irreproducible variability in subject 111312 include extensions beyond the group average in the retest but not test data for areas 45 and TPOJ3 (fuschia arrows in Fig 10D). Further inspection of Fig 10D-F reveals additional examples of both reproducible and irreproducible variability, but reproducible variability substantially outweighs the irreproducible variability for these 10 areas in these three subjects as expected.

### 3.7 Atypical area 55b organization

Area 55b, which is associated with language function, exhibits both typical and atypical patterns. The typical 55b, as seen in the group parcellation mentioned earlier, is situated between the frontal eye field (FEF) and premotor eye field (PEF) and is adjacent to M1 (area 4). However, atypical patterns have been reported, including: 1) Split 55b: In this pattern, 55b is split, resulting in FEF and PEF regions spatially adjoining (section #1: row 1 to 3 in Fig 11). 2)

**Figure 11.**
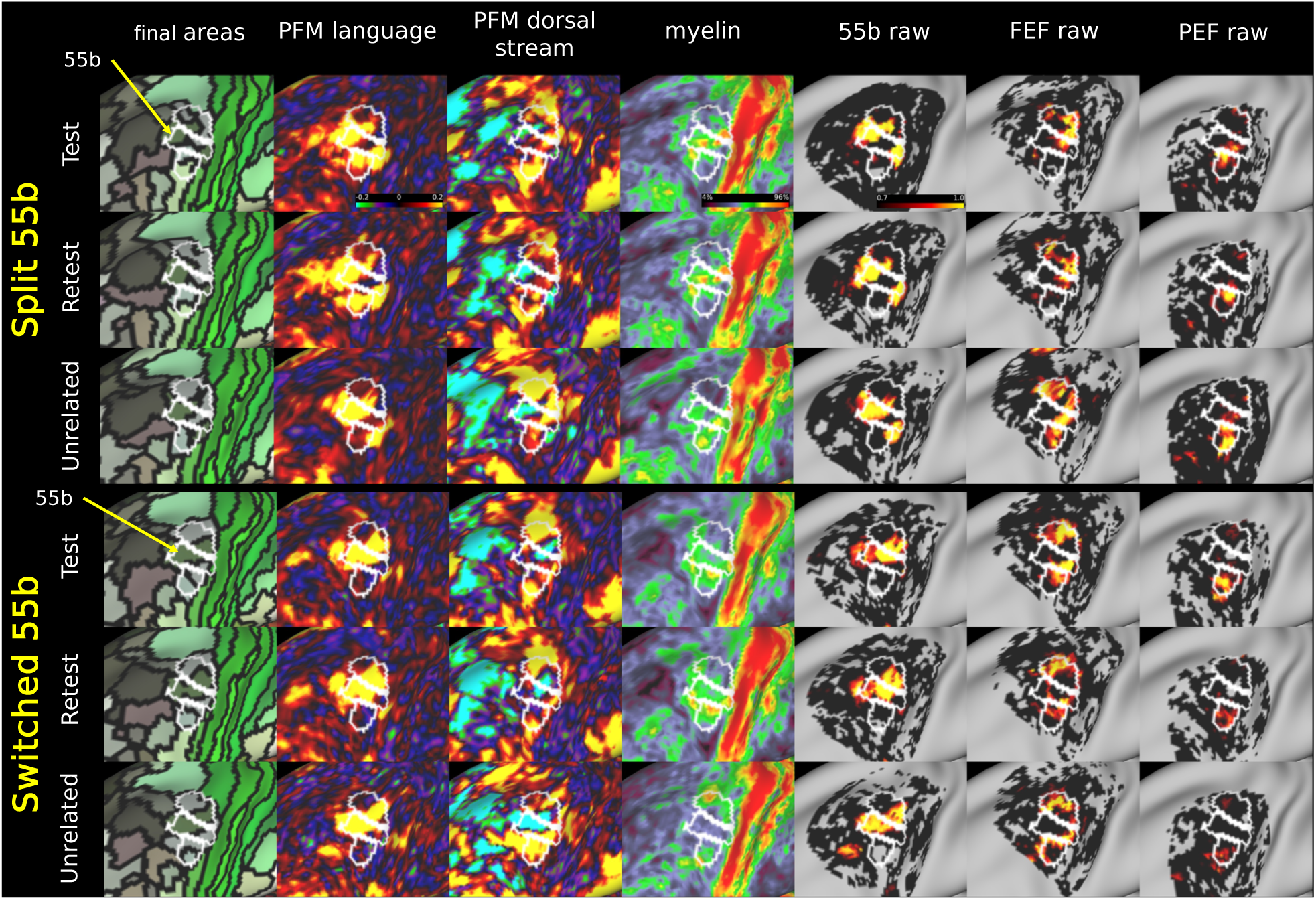
The atypical organizational patterns of area L_55b reported in the previous study (Glasser et al., 2016): Split (upper panels) and switched (lower panels) 55b. The individual parcellations for these atypical 55b organizations are depicted in column #1 for the same subjects in the test-retest groups and unrelated subjects in the 475V group who share the same 55b topology. Additionally, multimodal features, including the language network (column #2), the dorsal stream visual network (column #3), and myelin maps (column #4), are illustrated alongside the probabilistic maps for 55b, FEF, and PEF areas to provide a comprehensive view of these atypical patterns (the regressor output is clipped to 0 for predictions lower than 0). The areal boundaries in white are the L_55b, L_FEF and L_PEF in HCP_MMP1.0.

Switched (previously called shifted) 55b: In this atypical topology, 55b tilts upwards towards the superior-anterior portion of the upper limb somatosensory motor subregion, deviating from its typical ventral orientation (section #2: row 4 to 6) because it has switched positions with FEF, which again adjoins PEF. The language RSN map from PFM-WR is a strong indicator of typical vs atypical topology of 55b and the distinction between split and switched 55b. In addition to these known atypical patterns, this study identified two additional atypical subtypes:

1. Anterior 55b (section #3: row 1 to 3 in Fig 12): This variant is characterized by 55b not being adjacent to M1. It occurs when the language RSN map shows smaller or tilted activation in the typical 55b area and the dorsal stream visual RSN exhibits a horseshoe-shaped correlation peak adjacent to M1, but surrounding the language RSN in the typical 55b region.
2. Posterior 55b (section #4: row 4 to 6): This pattern is the opposite of anterior 55b, with the language RSN showing a 55b-related correlation peak adjacent to M1 in the 55b region, surrounded by a horseshoe-shaped dorsal stream correlation strip attached to 8Av.

**Figure 12.**
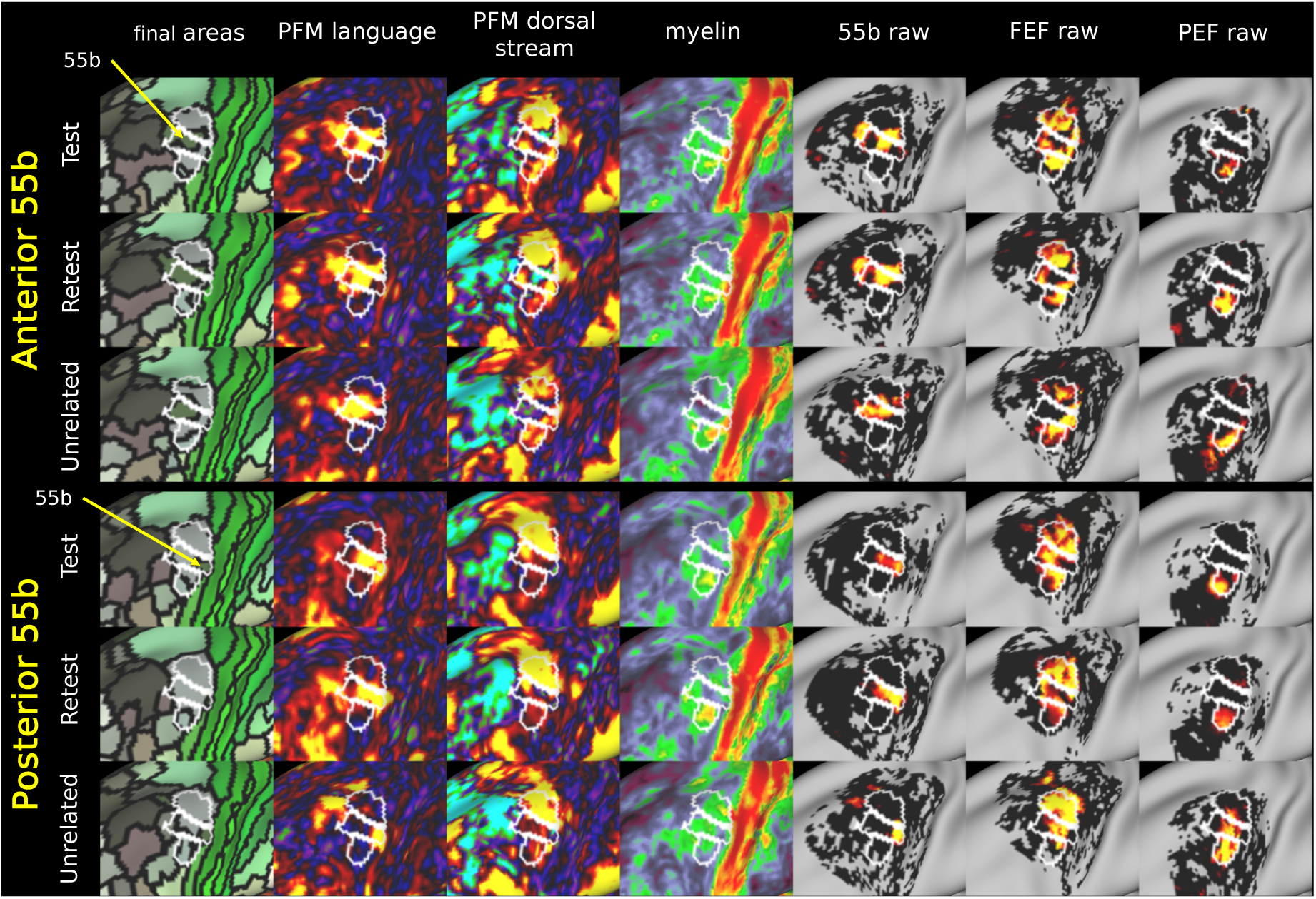
The newly reported atypical organizations of Area L_55b identified in this study: Anterior (upper panels) and posterior (lower panels).

Fig 11, Fig 12 and Fig S4 (zoom out) provide examples of these 55b atypical topologies predicted by the ARENA-2 model under the PFM-WR RSN setting. These findings underscore the multimodal nature of individual differences in cortical functional and structural organization, challenging the notion that these variations are mere noise in the data.

As in the original study (Glasser et al., 2016), we focused on atypical area 55b patterns in the left hemisphere (L_55b); atypicalities are also evident in the right hemisphere (R_55b) but have yet to be systematically examined. Manual inspection of all 1071 subjects revealed that 19.1% of the group exhibited atypical L_55b organization, 18.4% exhibited atypical R_55b organization (Table 4). Overall, counting atypical organization in both hemispheres, 30.2% of the group show atypical organization of area 55b in at least one hemisphere. That the incidence of atypicality in both hemispheres (7.4%) is only about twice that predicted by chance (0.191 x 0.184 ≈ 0.035 = 3.5%) suggests that area 55b atypicality is not simply a reflection of a genetic variant that has equivalent effects in the two hemispheres.

**Table 4.**
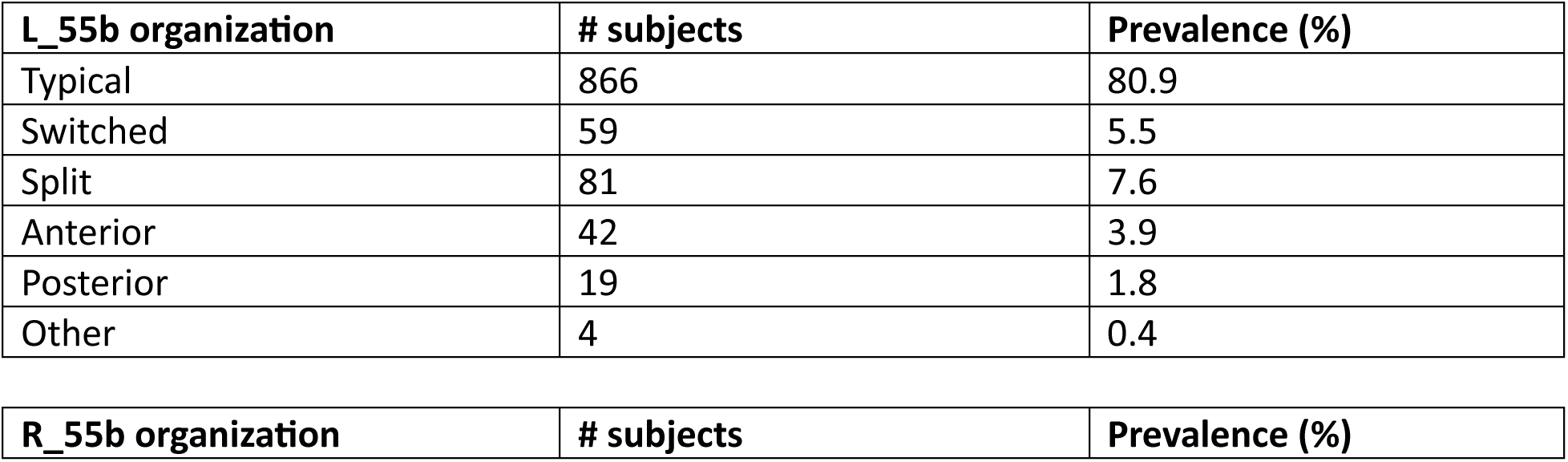
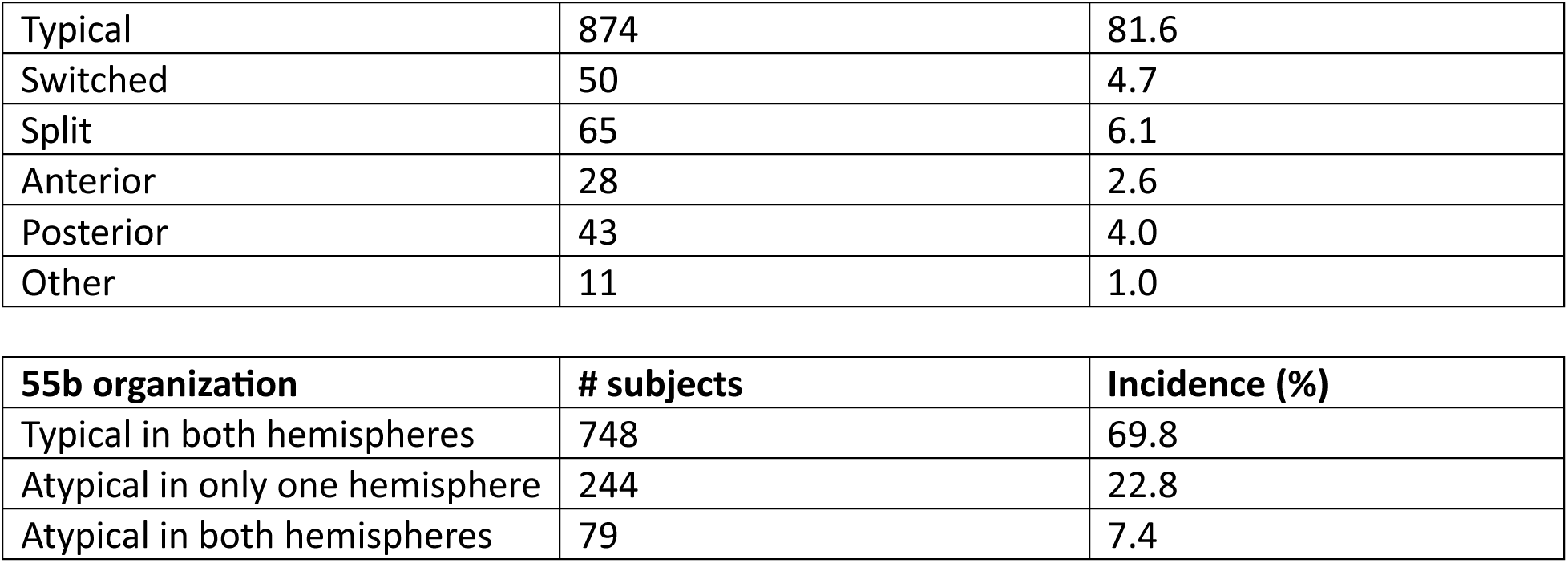
A summary of the typical and atypical organization for area L_55b and R_55b using the HCP-YA 1071 3T subjects with 1hr resting state fMRI and the PFM-WR d=76 approach.

| **L_55b organization** | **# subjects** | **Prevalence (%)** |
| --- | --- | --- |
| Typical | 866 | 80.9 |
| Switched | 59 | 5.5 |
| Split | 81 | 7.6 |
| Anterior | 42 | 3.9 |
| Posterior | 19 | 1.8 |
| Other | 4 | 0.4 |

| **R_55b organization** | **# subjects** | **Prevalence (%)** |
| --- | --- | --- |
| Typical | 874 | 81.6 |
| Switched | 50 | 4.7 |
| Split | 65 | 6.1 |
| Anterior | 28 | 2.6 |
| Posterior | 43 | 4.0 |
| Other | 11 | 1.0 |

| 55b organization | # subjects | Incidence (%) |
| --- | --- | --- |
| Typical in both hemispheres | 748 | 69.8 |
| Atypical in only one hemisphere | 244 | 22.8 |
| Atypical in both hemispheres | 79 | 7.4 |

## 4. Discussion

### 4.1 The different types of RSN maps

In the multimodal parcellation process, both structural and functional features are combined to create a comprehensive cortical map. However, functional features such as RSN maps can be decomposed in different ways, and which approach would be best for an areal classifier was unknown. Here, we investigated RSN maps generated by different methods, including spatial and temporal Independent Component Analysis (sICA and tICA), and PFM maps based on dual or weighted regression. We found that PFM-WR provides a balanced approach between classification performance and generalizability to new individuals and types of data.

Spatial ICA and temporal ICA involve a two-step procedure: first, estimating a group-level decomposition using ICA and then separately mapping these components onto individual subjects using either weighted spatial regression (for spatial ICA; (Glasser et al., 2016)) or single temporal regression (for temporal ICA; (Yang et al., 2024). Temporal ICA can also be applied to new individuals using weighted spatial regression first of the original sICA decomposition and then applying the tICA mixing matrix. PROFUMO, on the other hand, uses an iterative approach based on Bayesian decomposition that optimizes subject and group estimates, resulting in better subject-specific spatial variability compared to ICA. In our study (Results sections 3.2, 3.3 and 3.4), we made two key observations: 1) tICA consistently outperformed sICA RSNs in terms of reproducible individual variability (measured by RIV) and generalizability to an unseen fMRI paradigm. This is because tICA spatial maps contain all of the available connectivity information, whereas sICA retains some of the connectivity as correlations between component timeseries, leading to a reduction in information available to the areal classifier. (Temporal ICA decomposes the group fMRI matrix into a set of temporally independent timeseries and a set of correlated spatial maps, whereas spatial ICA decomposes the group fMRI matrix into a set of spatially independent spatial maps and a set of correlated timeseries). We found that tICA maps also resulted in better average areal detection rates and test-retest reproducibilities compared to sICA. 2) The original noise-free PROFUMO maps exhibited nearly perfect memorization of the training labels, resulting in exceptionally high average areal detection rates and test-retest reproducibility (Fig 6). However, this came at the cost of missing the more subtle features related to individual variability in noise-free PFM maps. We discovered that the high-contrast nature of the PFM origmaps made it easy for the classifier to learn where the areas were rather than their fingerprints based on area-specific features. For example, certain visual network maps in the PFM origmaps dataset were found to strongly influence prediction of the language area 55b (Refer to Fig S1) using the SHAP value (Lundberg & Lee, 2017) to explain the model’s decision process. In essence, the classifier learned that, for example, to fill in the gap in a split 55b caused by the eye fields coming together, it needed to use the dorsal stream visual network, rather than learning the genuinely different areal topology in this location involving 55b, FEF, and PEF. We attempted to avoid this behavior using a two-stage training framework for the areal classifier, initially training it using all subjects in the training dataset and selecting the most important features for each area to filter out subjects with poor alignment. In the second stage, we trained the areal classifier using only the most important features and well-aligned subjects. While this approach reintroduced some individual variability, the method performed less well than sICA, tICA, or PFM DR maps at capturing individual variability (Table S1).

To address these issues with PROFUMO maps, we applied dual regression to generate PFM-DR RSN maps. This was slightly preferred over noisy PFM spatial maps in the context of our study, as it can allow fairer comparisons to individualized tICA/sICA methods (see Methods section for details). PFM-DR reduced the contrast of the PFM features and avoided the overfitting problems of noise-free PFM origmaps. PFM-DR was the best method for identifying reliable individual variability that differed from the group parcellation HCP_MMP1.0_210P_Orig, performing better than tICA. However, PFM-DR is not currently the optimal choice, because PROFUMO does not currently support a straightforward approach for utilizing precomputed estimations on unseen groups of subjects or on the same group of subjects with different types of fMRI runs. To overcome this limitation, we developed a variant of PFM that uses weighted regression (PFM-WR), a fundamental step in the ICA-based methods. This approach allows the precomputed PROFUMO group decomposition to be applied to new individuals and new timeseries, yielding PFM-WR spatial maps and corresponding timeseries. We demonstrated that PFM-WR outperforms tICA in terms of reliable individual variability, averaged areal detection rate, and test-retest reproducibility, making it our currently preferred choice for areal classification.

However, as we plan to further develop the PFM technique to enable rapid estimation on new subjects or data types, PFM-DR could ultimately replace PFM-WR as our preferred RSN decomposition method for areal classification.

Another observation is that lower RSN dimensionality (d=76) estimated after six Wishart Distributions better captures reliable individual variability, not only for ICA-based RSN maps, sICA and tICA, but also for PFM-DR and PFM-WR (Table 3), with slightly worse averaged areal detection rate and test-retest reproducibility. This finding is likely because d=92 with WD=5 results in oversplitting of the resting state networks and inclusion of ‘unstructured’ noise (largely the most spatially autocorrelated noise), making it easier for the areal classifier to overfit and memorize the training labels. We also found that six Wishart distributions was the optimal number for temporal ICA cleanup (Yang et al., 2024).

### 4.2 Weakly supervised learning for areal classification

Conventional supervised learning for binary classification assumes that all labels are reliable, which is the approach adopted by the original MLP model for individual parcellation after eliminating subjects likely to be poorly matched to the group during an initial classification stage. However, due to individual variability in the features, assuming strict labels for learning can lead to suboptimal performance (Results Section 3.3), even when a filtering process is used to train only on well-aligned subjects based on the similarity between individual feature vectors and the group-averaged feature vector. Deriving individual parcellations from a group parcellation can be categorized as a weakly supervised learning problem, involving inaccurate supervision, where the group parcellation used as training labels (in-area or outside-area labels) should include uncertainty for every subject due to individual variability.

To our knowledge, this study represents the first application of weakly supervised learning to cortical parcellation, resulting in a refined areal classifier that achieves superior performance in every aspect when compared under the same RSN decompositions. We apply a two-stage strategy:

1. Ensemble learning is a simple yet effective strategy, where each base learner learns partial labels for a given area, and their predictions are aggregated to derive an initial prediction. This diversity is designed to mitigate the impact of label uncertainty. While some base learners may learn incorrect features or make errors on certain subsets of the data, the majority learn the right features and make correct predictions, thus reducing the overall noise in the ensemble.
2. A second-stage training uses a regression model to learn from the probabilities predicted in the first stage. This stage further mitigates label uncertainty and enhances the model’s capability to capture reproducible individual variability as demonstrated in Table 3. The rationale behind this approach is straightforward: by training the model to probabilistically learn the distribution of predictions using improved targets beyond binary group parcellation labels, we further enhance its performance. A variant of this method that we explored but did not use involves training the second stage using pseudo labels predicted in the first stage.

Notably, this multi-stage strategy may reduce training efficacy for some areas, potentially making them harder to identify. Validation against the initial group-level area size is necessary to address this issue. Supplementary Table S2 and Fig S6 demonstrate that two-stage training using pseudo labels significantly improves the measure of reliable individual variability.

However, this approach may also result in substantial changes in the mean size of some areas at the group MPM level compared to HCP_MMP1.0_210P_Orig, in contrast to the final approach of ARENA with pseudo probabilities, which allows the model to not only capture the predicted class, but also the uncertainty associated with each prediction. By considering uncertainty explicitly during training using pseudo probabilities, the model exhibits more stable behavior and produces less drastic changes in group mean areal size compared to using pseudo labels.

Further discussion about potential future directions utilizing weakly supervised learning is provided in section 4.4.

### 4.3 The individual parcellation framework

Alongside the refined areal classifier, we present a comprehensive individual parcellation framework designed for multimodal HCP-style datasets. This framework encompasses both an individual parcellation pipeline and guidelines for evaluating reproducible individual variability that differs from the group parcellation and generalizability to unseen fMRI paradigms. It serves as a reference for the development of future multimodal individual parcellation methods.

The framework begins with carefully cleaned fMRI data--including fMRI dense timeseries after sICA cleaning, recleaning, and tICA cleaning--to ensure that the derived areal organization is minimally influenced by artifacts, thereby improving the reliability of the resulting individual parcellations. The individual parcellation pipeline includes three main stages. Feature Generation: In this initial stage, maps of structural and functional features are generated for each subject. Inference by Areal Classifier: This key stage involves using pre-trained areal classifiers to assess the probability of the area class at each vertex within each area’s searchlight of the 360 areas defined in HCP_MMP1.0 in each individual subject. Individual Regularization: Several regularization steps are used to eliminate or merge small and isolated patches to create an individual’s parcellation. After the individual pipeline, group maps are generated with two additional stages. Combining Across Individuals: The individual-subject maps are combined across individuals to generate group probabilistic maps. Combination Across Probabilistic Areas: The final group-average cortex parcellation is obtained by combining across the probabilistic maps to generate a maximum probability map.

After obtaining the individual parcellations, the reliability of individual variability focuses on factors like avoiding overfitting to group training labels and maximizing the reproducible individual variability, such as the atypical topology of area 55b. The reliability can be quantified using a metric specifically designed for a group of test-retest subjects: Reliable Individual Variability, based on the individual Reliable Variability Score (iRVS), which reflects the ability of a model to capture reliable individual variability that differs from the group parcellation.

Another important component of evaluating an areal classifier is its generalizability to unseen fMRI paradigms, again focusing on reliable individual variability. The previously designed measures (average areal detection rate and test-retest reproducibility) can be manipulated by a model that perfectly learns to map the group training labels. We provide an example using PROFUMO origmaps (Fig 6), which can easily achieve nearly perfect average areal detection rate and test-retest reproducibility by overfitting to the training labels, but fails to capture reproducible individual variability, which is the primary purpose of an areal classifier in the setting of data that are reasonably well aligned.

In summary, the ARENA framework presented here is highly modular, allowing for updates to the group parcellation (labels), areal classifier, and types of multimodal features to adapt to evolving research needs. Additionally, it provides an unbiased evaluation strategy for any areal classifier that aims to capture reliable individual variability. Its versatility makes it a tool for the further refinement of multimodal-based individual parcellation methods.

### 4.4 Limitations and future work

This work leaves several important questions unanswered. For instance, it remains to be determined how the performance of the areal classifier depends on various data acquisition parameters. The total duration of fMRI scans is of particular interest, because most studies acquire much less than the nearly 60 min (4 x ∼15 min) acquired from HCP-YA subjects.

Additionally, it is unknown how classifier performance would change with even more data or how it would perform on 7T datasets. More generally, it is unknown whether areal classifier performance is currently limited primarily by SNR efficiency or total acquisition time of fMRI (i.e., whether better sampling of a given amount of time is needed or whether it is more important to sample a wider range of brains states by sampling for longer). Generalizability to different age ranges or disease states is also unknown, as is the question of whether there are any new insights into areal organization, such as the discovery of missing or extra areas in the 1071 HCP-YA subjects. We have primarily focused on developing methodology for individual parcellation, and these questions warrant exploration as part of future work. Given its superior robustness, reproducibility, and reliability, we anticipate that this refined areal classifier will permit new neurobiological findings such as the diversity of topologies of area 55b.

Besides ensemble learning and dual-stage training schema, other directions that might benefit from a weak supervised learning approach are left for future work. 1) Label re-design based on spatial uncertainty during the first stage: As the group parcellation does not represent variability at the individual level, the in-area label (value equal to 1) or the outside-area label (value equal to 0) can be adjusted to have intermediate values based on how close to the edge of the original binary label locations are. In this case, the learning process may be stabilized by not being given a hard edge to try to overfit, thus increasing the model’s generalizability. 2) Loss function design: The current classification model doesn’t consider the cortical surface areas associated with each vertex when making the predictions, which could be improved by a customized loss function design. 3) Use of spatially aware machine learning methods: Deep learning might permit elimination of the ad hoc regularization step at the end of the individual classification process. On the other hand, deep learning will be much more prone to simply learning the location of training labels, rather than their multi-modal fingerprints, and might not provide better reproducible individual variability.

A major current limitation is that the MSMAll areal feature-based surface registration used in this study is, for comparison reasons, not the most recent iteration of the MSM regularization model (the strain-based one of Robinson et al. (2018), but rather an older angular deviation model with higher order regularization (Glasser et al., 2016; Robinson et al., 2014). Additionally, even the latest version of MSMAll (used for HCP Lifespan) has not been optimized for test-retest reproducibility. Thus, because test-retest reproducibility of the areal classifier is evaluated through the MSMAll alignment to a common atlas, any errors in either test or retest in the registration alignment will penalize the areal classifier. For example, in Figure 4B it appears that the anterior split portion of 55b has been shifted anteriorly and superiorly by about 1 vertex in the test relative to the retest dataset. Such errors would affect all areal classifier candidates equivalently and thus are not relevant to determining the best areal classifier; however, when the registration is improved, the reproducibility-based performance estimates presented here will improve without a change to the classifier.

## 5. Conclusions

We have demonstrated a refined multimodal individual parcellation method that yields reliable measures of subject-specific cortical areal variability differing from the group parcellation and strong generalizability to unseen fMRI types. The superior performance is achieved by 1) An ensemble classifier with dual training stages that treats the problem as a type of weakly supervised learning. 2) Highly reproducible PROFUMO RSN maps generated after weighted regression that can easily be applied to unseen group or fMRI types. We introduced a modular individual parcellation framework to enable plug-and-play usage of any future-developed advanced areal classifier. To thoroughly evaluate the areal classifier, we designed new metrics that can quantify ability to capture reproducible individual variability for each area at the individual and group level, and additionally evaluated its generalizability to unseen fMRI types, paving the foundation for future studies. We generated a comprehensive summary of the organization of area 55b in the left and right hemispheres using HCP-Young Adult 3T 1071 subjects, along with identifying new atypical 55b topologies: anterior 55b and posterior 55b.

The automated individual parcellation pipeline has been integrated into the HCP pipelines, allowing users to generate subject-specific parcellations easily. We will release the individual parcellation data for the HCP-Young Adult 1071 subjects simultaneously.

## Acknowledgments

Data were provided in part by HCP-YA: the Human Connectome Project, WU-Minn Consortium (Principal Investigators: David Van Essen and Kamil Ugurbil; 1U54MH091657) funded by the 16 NIH Institutes and Centers that support the NIH Blueprint for Neuroscience Research and by the McDonnell Center for Systems Neuroscience at Washington University. Supported in part by Connectome Coordination Facility (CCF) I: R24MH108315, Connectome Coordination Facility II: 1R24MH122820, the McDonnell Center for Systems Neuroscience at Washington University, and NIH R01MH060974 (DCVE, MFG). Computations were performed using the facilities of the Washington University Research Computing and Informatics Facility, which were partially funded by NIH grants S10OD025200, 1S10RR022984-01A1 and 1S10OD018091-01. Additional support was provided by The McDonnell Center for Systems Neuroscience.

We thank Emma Robinson and Logan Williams from King’s College London for valuable discussions. We thank Jennifer S. Elam for help with editing the initial draft of the manuscript for English Grammar and Style.

## Data and Code Availability

The datasets used in this study are available in ConnectomeDB (https://db.humanconnectome.org/app/template/Login.vm). The code for the HCP Pipelines is available in https://github.com/Washington-University/HCPpipelines.

## Author Contributions

C.Y. and M.F.G. conceptualized this study and designed the methodology. C.Y. conducted the data analysis, ran the experiments, and wrote the first draft. T.S.C., S.R.F., J.D.B., S.M.S., D.V.E. and M.F.G. contributed to the revisions to the manuscript. T.S.C., S.R.F., J.D.B., S.M.S., D.V.E. and M.F.G. provided guidance on data analysis and results interpretation. D.V.E. and M.F.G. supervised the work.

## Supplementary Figures and Tables

**Supplementary Figure 1.**
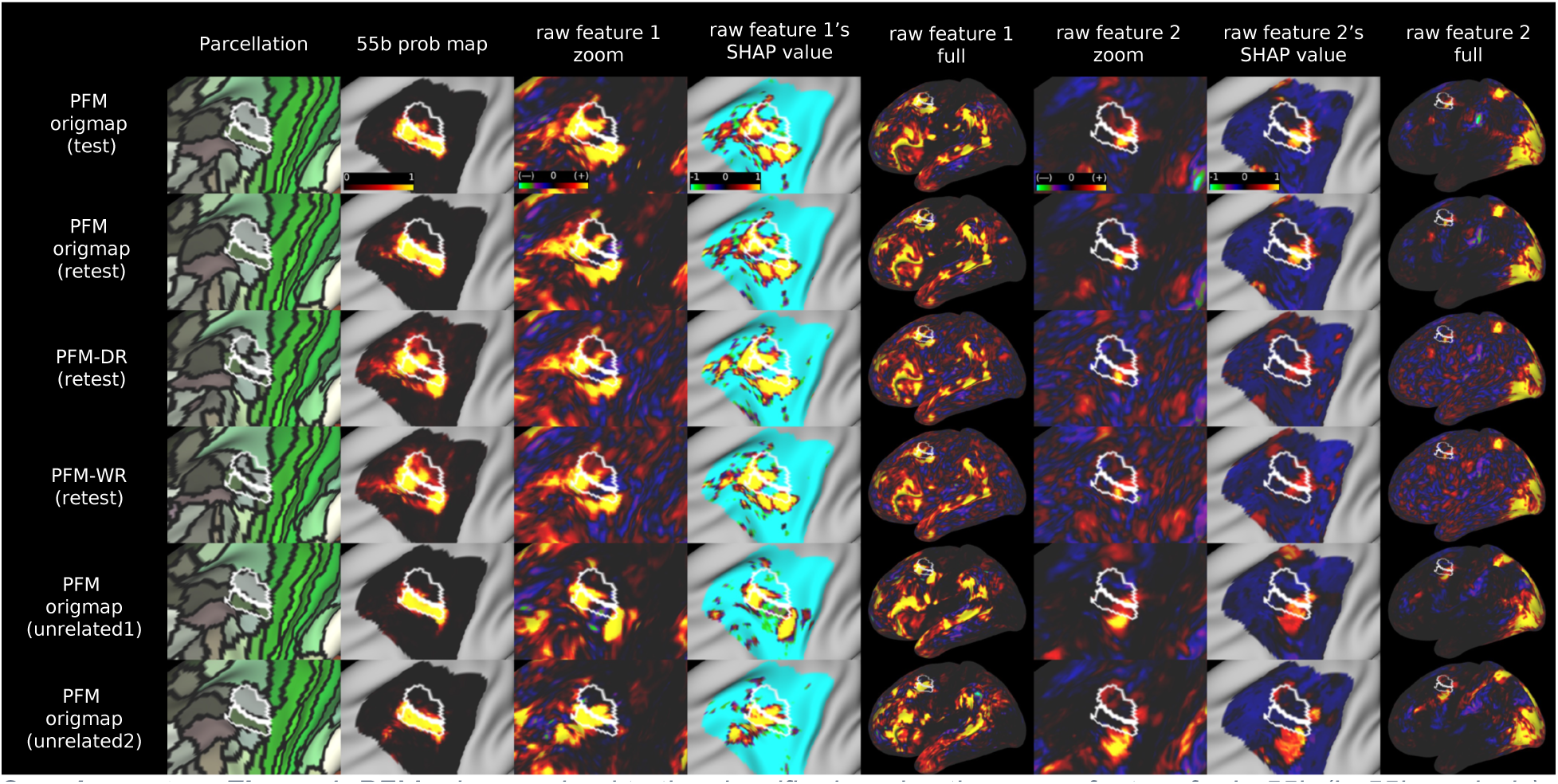
PFM origmaps lead to the classifier learning the wrong feature for L_55b (L_55b analysis). From top to bottom, the predicted individual parcellations are illustrated for the same test-retest subject (149337) with atypical 55b in the left hemisphere (rows 1-4) and two unrelated subjects (153227 and 100206) in the 475V evaluation group (split 55b in row 5 and switched 55b in row 6). The SHAP value (columns 5 and 7) scaled from −1 to 1 is computed for the feature maps in the search region of L_55b to represent the importance of the feature when the model makes the final prediction. The language network feature map (columns 3 and 5) clearly shows a split 55b topology indicating strong evidence of a split 55b areal organization, however, the probability maps (rows 1, 2 and 5) strongly suggest a typical organization. This is due to an incorrect feature learned by the model to determine the 55b prediction: a visual network feature map (rows 1, 2 and 5 under columns 6-8) that has a high SHAP value to fill in the gap between the split 55b region. Moreover, PFM-DR (row 3 column 6-8) and PFM-WR (row 4 column 6-8) with noise reintroduced back has a reduced contribution from the same visual network map.

**Supplementary Figure 2.**
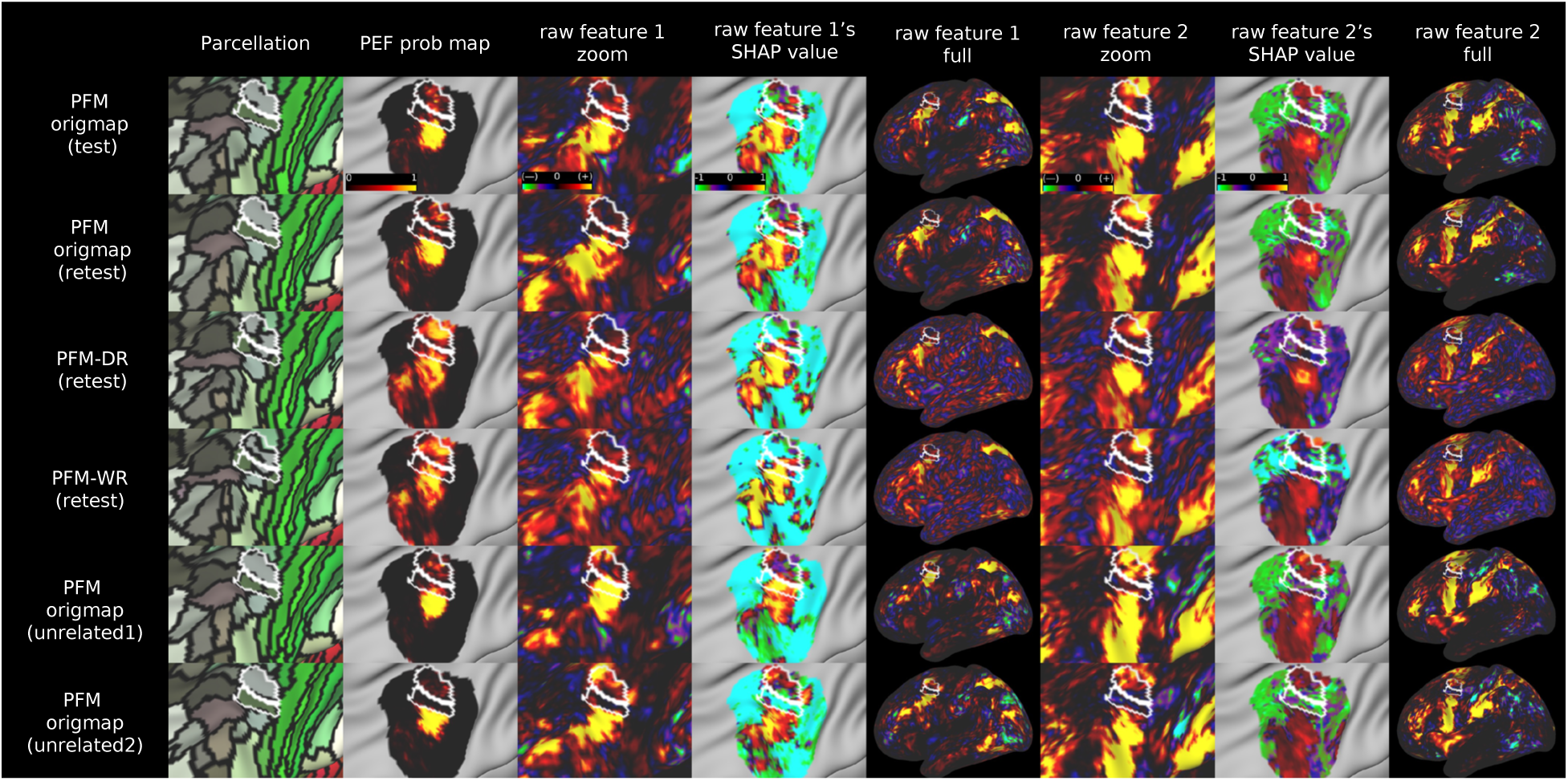
PFM origmaps leads to the classifier learning a wrong feature for L_55b (L_PEF analysis).

**Supplementary Figure 3.**
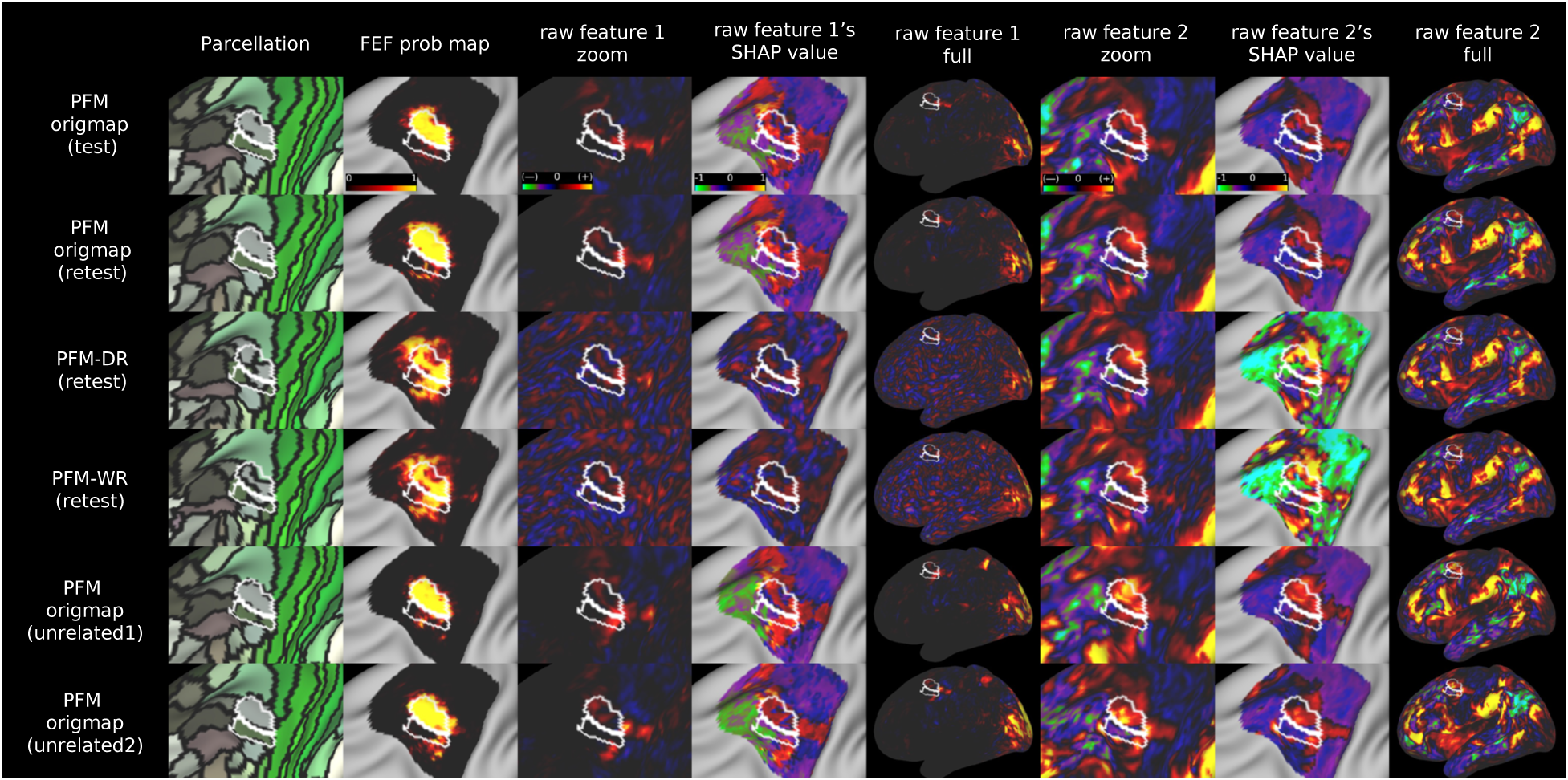
PFM origmaps leads to the classifier learning a wrong feature for L_55b (L_FEF analysis).

**Supplementary Figure 4.**
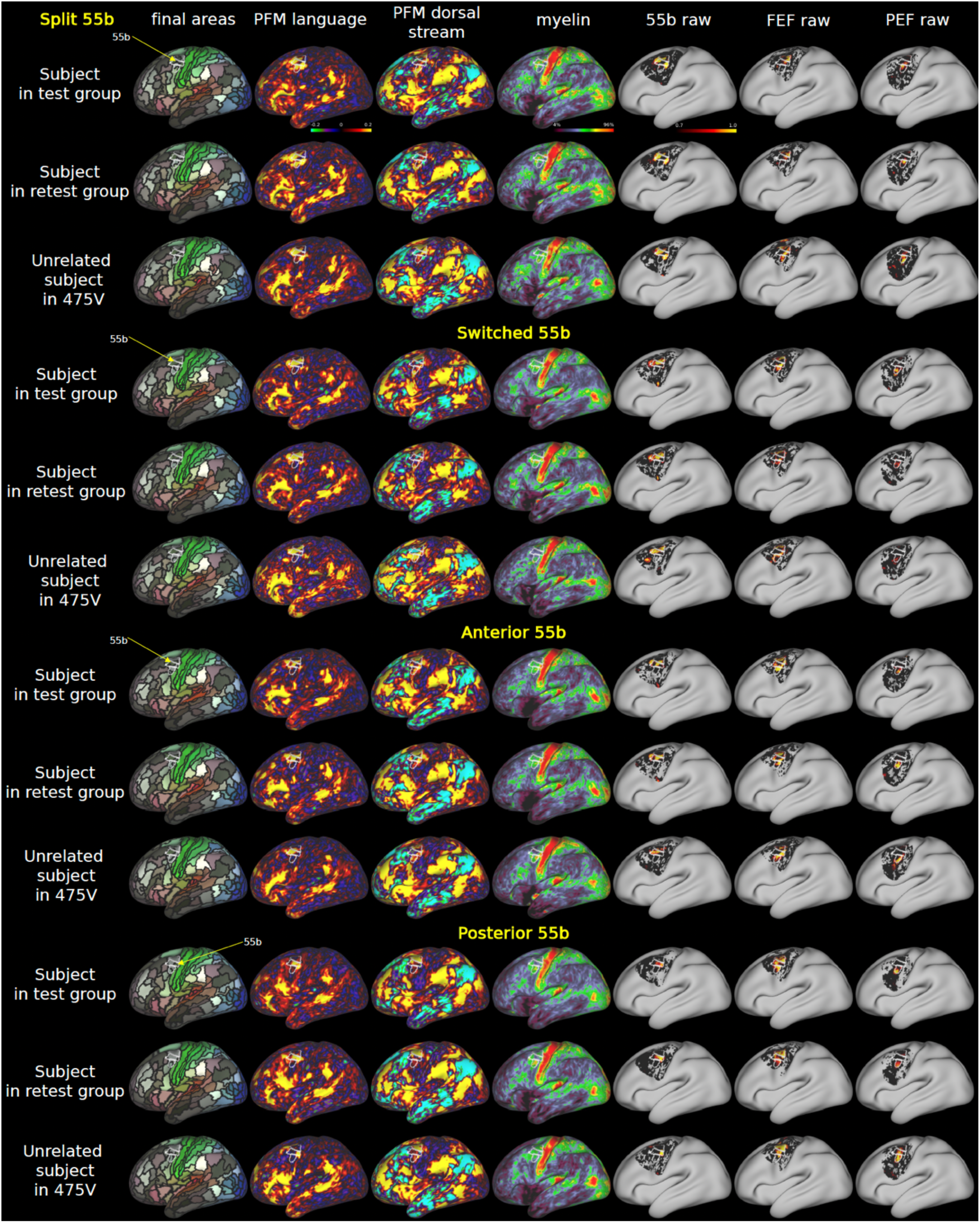
Exemplars for atypical 55b organization in the left hemisphere (zoomed out version of main Figure 10 and 11).

**Supplementary Figure 5.**
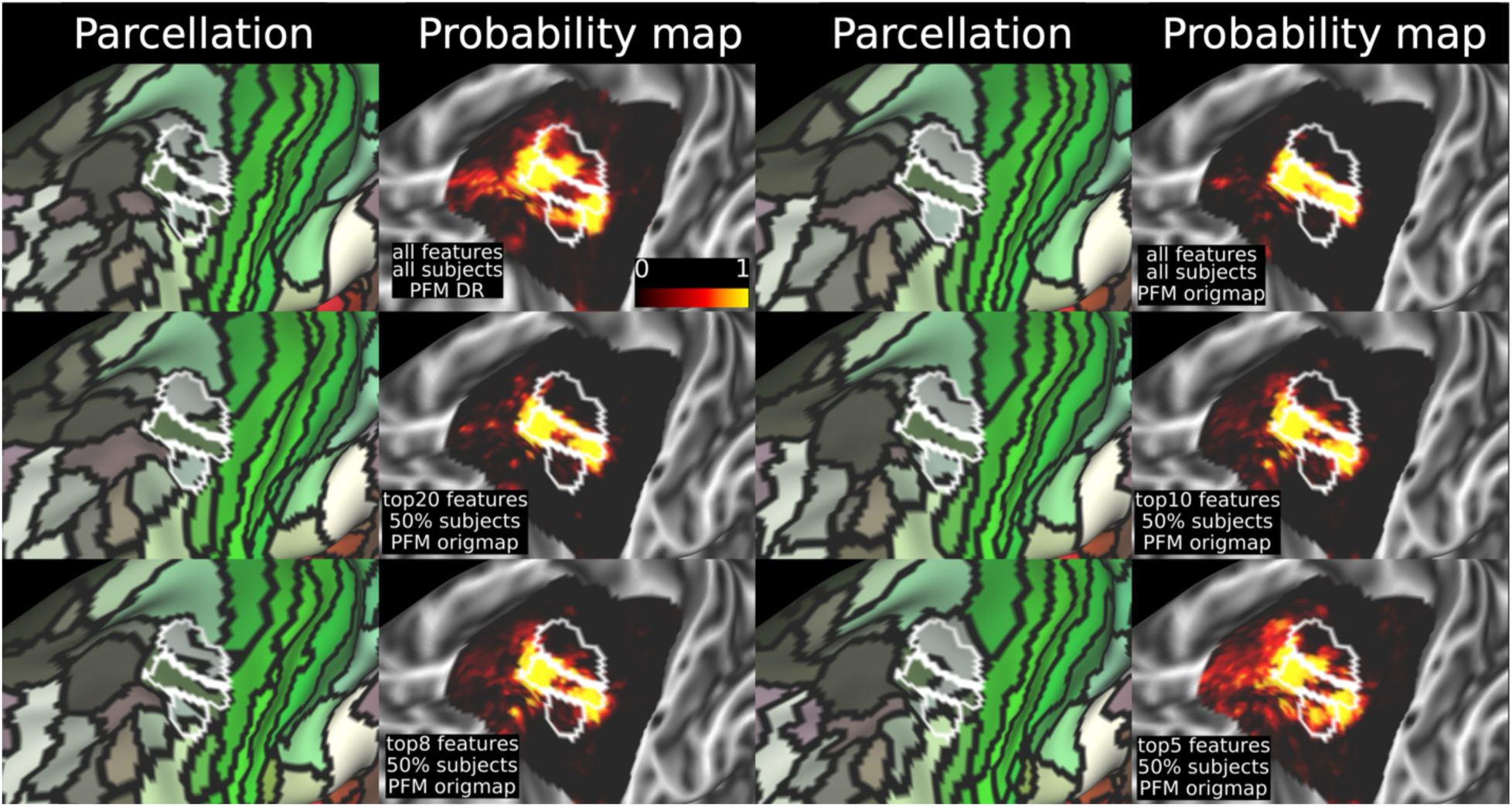
The individual variability of 55b can be predicted after a strict feature selection under the RSN decomposition of PFM origmaps. All the parcellations are predicted on the same retest subject which has split 55b in the language network RSN map.

**Supplementary Figure 6.**
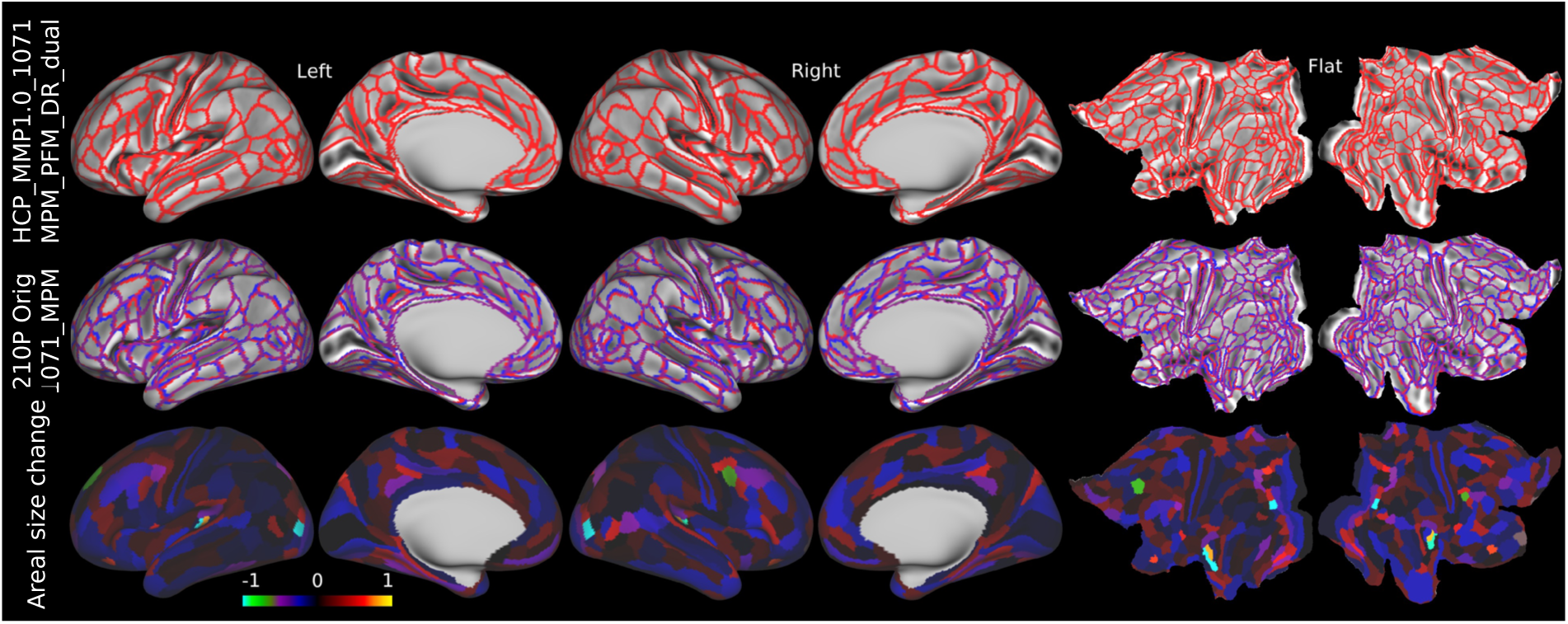
Dual training stage with PFM-DR increases the model’s ability to capture reproducible individual variability but also increases the deviation from the original group parcellation. Compared to main Figure 9, more areas have substantially changed area size compared to HCP_MMP1.0_210P_Orig (row #3). The Dice overlap between the original semi-automated parcellation (HCP_MMP1.0_210P_Orig) and the 1071 group MPM under PFM-DR with dual stages (trained on pseudo labels) was 0.867 (versus 0.880 for ARENA-2; trained on pseudo probabilities). In comparison, the Dice overlap between the original semi-automated parcellation and the 1071 group MPM under PFM-WR with dual stages was 0.870 (versus 0.888 for ARENA-2 shown in main Figure 9).

**Supplementary Table 1.** Performance evaluation with PFM origmaps using feature and subject selections. The top three rows are the three ARENA-1 areal classifiers with PFM-WR, PFM-DR and tICA d=76. (replicated from main Table 3)

| Training parameters | Score-based Reliable Individual variability | Test-retest reproducibility (%) | Averaged areal detection rate (%) |
| --- | --- | --- | --- |
| PFM-WR d=76 | 7416 | 73.8 | 98.10 |
| PFM-DR d=76 | 8876 | 72.6 | 97.54 |
| tICA d=76 | 7276 | 73.3 | 97.91 |
| PFM origmaps below d=76 |  |  |  |
| Raw (all features and all subjects) | -1998 | 86.9 | 99.94 |
| Top20 features top50% well-aligned subjects | -1236 | 81.5 | 99.38 |
| Top10 features top50% well-aligned subjects | -246 | 75.8 | 98.21 |
| Top8 features top50% well-aligned subjects | 736 | 73.3 | 97.77 |
| Top5 features top50% well-aligned subjects | 2864 | 67.2 | 95.09 |

**Supplementary Table 2.** Performance evaluation under two-stage training where the 2^nd^ stage uses the predicted area labels from the 1^st^ stage as training targets. This strategy can substantially improve the mRIV, showing a better ability to capture reproducible individual variability differing from the group parcellation. The top three rows are the three best areal classifiers with only one-stage training (ARENA-1). (replicated from main Table 3). The top performer is in bold.

| Setting<br>(d=76) | median Reliable<br>Individual variability | Test-retest<br>reproducibility (%) | Averaged areal<br>detection rate (%) |
| --- | --- | --- | --- |
| PFM-WR (ARENA-1) | 7416 | 73.8 | 98.10 |
| PFM-DR (ARENA-1) | 8876 | 72.6 | 97.54 |
| PFM-WR (ARENA-2) | 8380 | 73.3 | 97.81 |
| PFM-DR (ARENA-2) | 9786 | 71.5 | 97.28 |
| PFM-WR two stages<br>(2<sup>nd</sup> stage use individual<br>predicted areas as labels) | 10438 | 72.3 | 97.49 |
| PFM-DR two stages<br>(2<sup>nd</sup> stage use individual<br>predicted areas as labels) | **10448** | 72.1 | 97.27 |

## References

Andrews, T. J., Halpern, S. D., & Purves, D. (1997). Correlated size variations in human visual cortex, lateral geniculate nucleus, and optic tract. Journal of neuroscience, 17(8), 2859–2868.

Barch, D. M., Burgess, G. C., Harms, M. P., Petersen, S. E., Schlaggar, B. L., Corbetta, M., Glasser, M. F., Curtiss, S., Dixit, S., Feldt, C., Nolan, D., Bryant, E., Hartley, T., Footer, O., Bjork, J. M., Poldrack, R., Smith, S., Johansen-Berg, H., Snyder, A. Z., & Van Essen, D. C. (2013). Function in the human connectome: task-fMRI and individual differences in behavior. Neuroimage, 80, 169–189. 10.1016/j.neuroimage.2013.05.033

Bijsterbosch, J. D., Woolrich, M. W., Glasser, M. F., Robinson, E. C., Beckmann, C. F., Van Essen, D. C., Harrison, S. J., & Smith, S. M. (2018). The relationship between spatial configuration and functional connectivity of brain regions. Elife, 7. 10.7554/eLife.32992

Brodmann, K. (1909). Vergleichende Lokalisationslehre der Grosshirnrinde in ihren Prinzipien dargestellt auf Grund des Zellenbaues. Johann Ambrosius Barth.

Cachia, A., Roell, M., Mangin, J. F., Sun, Z. Y., Jobert, A., Braga, L., Houde, O., Dehaene, S., & Borst, G. (2018). How interindividual differences in brain anatomy shape reading accuracy. Brain Struct Funct, 223(2), 701–712. 10.1007/s00429-017-1516-x

Chen, T., & Guestrin, C. (2016). Xgboost: A scalable tree boosting system. Proceedings of the 22nd acm sigkdd international conference on knowledge discovery and data mining.

Claesen, M., De Smet, F., Suykens, J. A. K., & De Moor, B. (2015). A robust ensemble approach to learn from positive and unlabeled data using SVM base models. Neurocomputing, 160, 73–84. 10.1016/j.neucom.2014.10.081

Farahibozorg, S.-R., Bijsterbosch, J. D., Gong, W., Jbabdi, S., Smith, S. M., Harrison, S. J., & Woolrich, M. W. (2021). Hierarchical modelling of functional brain networks in population and individuals from big fMRI data. Neuroimage, 243, 118513. 10.1016/j.neuroimage.2021.118513

Fischl, B., Rajendran, N., Busa, E., Augustinack, J., Hinds, O., Yeo, B. T., Mohlberg, H., Amunts, K., & Zilles, K. (2008). Cortical folding patterns and predicting cytoarchitecture. Cerebral cortex, 18(8), 1973–1980.

Glasser, M. F., Coalson, T. S., Bijsterbosch, J. D., Harrison, S. J., Harms, M. P., Anticevic, A., Van Essen, D. C., & Smith, S. M. (2018). Using temporal ICA to selectively remove global noise while preserving global signal in functional MRI data. Neuroimage, 181, 692–717. 10.1016/j.neuroimage.2018.04.076

Glasser, M. F., Coalson, T. S., Bijsterbosch, J. D., Harrison, S. J., Harms, M. P., Anticevic, A., Van Essen, D. C., & Smith, S. M. (2019). Classification of temporal ICA components for separating global noise from fMRI data: Reply to Power. Neuroimage, 197, 435–438. 10.1016/j.neuroimage.2019.04.046

Glasser, M. F., Coalson, T. S., Harms, M. P., Xu, J., Baum, G. L., Autio, J. A., Auerbach, E. J., Greve, D. N., Yacoub, E., Van Essen, D. C., Bock, N. A., & Hayashi, T. (2022). Empirical transmit field bias correction of T1w/T2w myelin maps. Neuroimage, 258, 119360. 10.1016/j.neuroimage.2022.119360

Glasser, M. F., Coalson, T. S., Robinson, E. C., Hacker, C. D., Harwell, J., Yacoub, E., Ugurbil, K., Andersson, J., Beckmann, C. F., Jenkinson, M., Smith, S. M., & Van Essen, D. C. (2016). A multi-modal parcellation of human cerebral cortex. Nature, 536(7615), 171–178. 10.1038/nature18933

Glasser, M. F., Sotiropoulos, S. N., Wilson, J. A., Coalson, T. S., Fischl, B., Andersson, J. L., Xu, J., Jbabdi, S., Webster, M., Polimeni, J. R., Van Essen, D. C., Jenkinson, M., & Consortium, W. U.-M. H. (2013). The minimal preprocessing pipelines for the Human Connectome Project. Neuroimage, 80, 105–124. 10.1016/j.neuroimage.2013.04.127

Glasser, M. F., & Van Essen, D. C. (2011). Mapping human cortical areas in vivo based on myelin content as revealed by T1-and T2-weighted MRI. Journal of neuroscience, 31(32), 11597–11616.

Gordon, E. M., Laumann, T. O., Adeyemo, B., Huckins, J. F., Kelley, W. M., & Petersen, S. E. (2016). Generation and evaluation of a cortical area parcellation from resting-state correlations. Cerebral cortex, 26(1), 288–303.

Hacker, C. D., Laumann, T. O., Szrama, N. P., Baldassarre, A., Snyder, A. Z., Leuthardt, E. C., & Corbetta, M. (2013). Resting state network estimation in individual subjects. Neuroimage, 82, 616–633. 10.1016/j.neuroimage.2013.05.108

Harrison, S. J., Bijsterbosch, J. D., Segerdahl, A. R., Fitzgibbon, S. P., Farahibozorg, S.-R., Duff, E. P., Smith, S. M., & Woolrich, M. W. (2020). Modelling subject variability in the spatial and temporal characteristics of functional modes. Neuroimage, 222, 117226. 10.1016/j.neuroimage.2020.117226

Harrison, S. J., Woolrich, M. W., Robinson, E. C., Glasser, M. F., Beckmann, C. F., Jenkinson, M., & Smith, S. M. (2015). Large-scale probabilistic functional modes from resting state fMRI. Neuroimage, 109, 217–231. 10.1016/j.neuroimage.2015.01.013

Hilgetag, C. C., & Barbas, H. (2006). Role of mechanical factors in the morphology of the primate cerebral cortex. PLoS computational biology, 2(3), e22.

Høie, M., Gade, F., Johansen, J., Würtzen, C., Winther, O., Nielsen, M., Marcatili, P. (2023). DiscoTope-3.0 - Improved B-cell epitope prediction using AlphaFold2 modeling and inverse folding latent representations. bioRxiv, 2023.2002.2005.527174. 10.1101/2023.02.05.527174

Kong, R., Li, J., Orban, C., Sabuncu, M. R., Liu, H., Schaefer, A., Sun, N., Zuo, X. N., Holmes, A. J., Eickhoff, S. B., & Yeo, B. T. T. (2019). Spatial Topography of Individual-Specific Cortical Networks Predicts Human Cognition, Personality, and Emotion. Cereb Cortex, 29(6), 2533–2551. 10.1093/cercor/bhy123

Kong, R., Yang, Q., Gordon, E., Xue, A., Yan, X., Orban, C., Zuo, X. N., Spreng, N., Ge, T., Holmes, A., Eickhoff, S., & Yeo, B. T. T. (2021). Individual-Specific Areal-Level Parcellations Improve Functional Connectivity Prediction of Behavior. Cereb Cortex, 31(10), 4477–4500. 10.1093/cercor/bhab101

Lundberg, S. M., & Lee, S.-I. (2017). A unified approach to interpreting model predictions. Advances in neural information processing systems, 30.

Mordelet, F., & Vert, J. P. (2014). A bagging SVM to learn from positive and unlabeled examples. Pattern Recognition Letters, 37, 201–209. 10.1016/j.patrec.2013.06.010

Nieuwenhuys, R., Broere, C. A., & Cerliani, L. (2015). A new myeloarchitectonic map of the human neocortex based on data from the Vogt–Vogt school. Brain Structure and Function, 220, 2551–2573.

Power, J. D., Cohen, A. L., Nelson, S. M., Wig, G. S., Barnes, K. A., Church, J. A., Vogel, A. C., Laumann, T. O., Miezin, F. M., Schlaggar, B. L., & Petersen, S. E. (2011). Functional network organization of the human brain. Neuron, 72(4), 665–678. 10.1016/j.neuron.2011.09.006

Robinson, E. C., Garcia, K., Glasser, M. F., Chen, Z., Coalson, T. S., Makropoulos, A., Bozek, J., Wright, R., Schuh, A., Webster, M., Hutter, J., Price, A., Cordero Grande, L., Hughes, E., Tusor, N., Bayly, P. V., Van Essen, D. C., Smith, S. M., Edwards, A. D.,…Rueckert, D. (2018). Multimodal surface matching with higher-order smoothness constraints. Neuroimage, 167, 453–465. 10.1016/j.neuroimage.2017.10.037

Robinson, E. C., Jbabdi, S., Glasser, M. F., Andersson, J., Burgess, G. C., Harms, M. P., Smith, S. M., Van Essen, D. C., & Jenkinson, M. (2014). MSM: a new flexible framework for Multimodal Surface Matching. Neuroimage, 100, 414–426. 10.1016/j.neuroimage.2014.05.069

Salimi-Khorshidi, G., Douaud, G., Beckmann, C. F., Glasser, M. F., Griffanti, L., & Smith, S. M. (2014). Automatic denoising of functional MRI data: combining independent component analysis and hierarchical fusion of classifiers. Neuroimage, 90, 449–468. 10.1016/j.neuroimage.2013.11.046

Schaefer, A., Kong, R., Gordon, E. M., Laumann, T. O., Zuo, X. N., Holmes, A. J., Eickhoff, S. B., & Yeo, B. T. T. (2018). Local-Global Parcellation of the Human Cerebral Cortex from Intrinsic Functional Connectivity MRI. Cereb Cortex, 28(9), 3095–3114. 10.1093/cercor/bhx179

Sheng, V. S., Provost, F., & Ipeirotis, P. G. (2008). *Get another label? improving data quality and data mining using multiple, noisy labelers* Proceedings of the 14th ACM SIGKDD international conference on Knowledge discovery and data mining, Las Vegas, Nevada, USA. 10.1145/1401890.1401965

Smith, S. M., Beckmann, C. F., Andersson, J., Auerbach, E. J., Bijsterbosch, J., Douaud, G., Duff, E., Feinberg, D. A., Griffanti, L., Harms, M. P., Kelly, M., Laumann, T., Miller, K. L., Moeller, S., Petersen, S., Power, J., Salimi-Khorshidi, G., Snyder, A. Z., Vu, A. T.,…Consortium, W. U.-M. H. (2013). Resting-state fMRI in the Human Connectome Project. Neuroimage, 80, 144–168. 10.1016/j.neuroimage.2013.05.039

Snow, R., O’Connor, B., Jurafsky, D., & Ng, A. Y. (2008). Cheap and fast---but is it good? evaluating non-expert annotations for natural language tasks Proceedings of the Conference on Empirical Methods in Natural Language Processing, Honolulu, Hawaii.

Stensaas, S. S., Eddington, D. K., & Dobelle, W. H. (1974). The topography and variability of the primary visual cortex in man. Journal of neurosurgery, 40(6), 747–755.

Ugurbil, K., Xu, J., Auerbach, E. J., Moeller, S., Vu, A. T., Duarte-Carvajalino, J. M., Lenglet, C., Wu, X., Schmitter, S., Van de Moortele, P. F., Strupp, J., Sapiro, G., De Martino, F., Wang, D., Harel, N., Garwood, M., Chen, L., Feinberg, D. A., Smith, S. M.,…Consortium, W. U.-M. H. (2013). Pushing spatial and temporal resolution for functional and diffusion MRI in the Human Connectome Project. Neuroimage, 80, 80–104. 10.1016/j.neuroimage.2013.05.012

Van Essen, D. C. (1997). A tension-based theory of morphogenesis and compact wiring in the central nervous system. Nature, 385(6614), 313–318.

Van Essen, D. C., Donahue, C. J., Coalson, T. S., Kennedy, H., Hayashi, T., & Glasser, M. F. (2019). Cerebral cortical folding, parcellation, and connectivity in humans, nonhuman primates, and mice. Proceedings of the National Academy of Sciences, 116(52), 26173–26180.

Van Essen, D. C., & Glasser, M. F. (2018). Parcellating Cerebral Cortex: How Invasive Animal Studies Inform Noninvasive Mapmaking in Humans. Neuron, 99(4), 640–663. 10.1016/j.neuron.2018.07.002

Van Essen, D. C., Smith, S. M., Barch, D. M., Behrens, T. E., Yacoub, E., Ugurbil, K., & Consortium, W. U.-M. H. (2013). The WU-Minn Human Connectome Project: an overview. Neuroimage, 80, 62–79. 10.1016/j.neuroimage.2013.05.041

Vogt, C., & Vogt, O. (1919). Allgemeine ergebnisse unserer hirnforschung (Vol. 25). JA Barth.

Yang, C., Coalson, T. S., Smith, S. M., Elam, J. S., Essen, D. C. V., & Glasser, M. F. (2024). Automating the Human Connectome Project’s Temporal ICA Pipeline. bioRxiv, 2024.2001.2015.574667. 10.1101/2024.01.15.574667

Yeo, B. T., Krienen, F. M., Sepulcre, J., Sabuncu, M. R., Lashkari, D., Hollinshead, M., Roffman, J. L., Smoller, J. W., Zöllei, L., & Polimeni, J. R. (2011). The organization of the human cerebral cortex estimated by intrinsic functional connectivity. Journal of neurophysiology.

Zhao, Z., Zeng, Z., Xu, K., Chen, C., & Guan, C. (2021). DSAL: Deeply Supervised Active Learning From Strong and Weak Labelers for Biomedical Image Segmentation. IEEE J Biomed Health Inform, 25(10), 3744–3751. 10.1109/JBHI.2021.3052320

Zhou, Z.-H. (2012). Ensemble Methods: Foundations and Algorithms. Chapman & Hall/CRC.

Zhou, Z.-H. (2017). A brief introduction to weakly supervised learning. National Science Review, 5(1), 44-53. 10.1093/nsr/nwx106

